# Early or late gestational exposure to maternal immune activation alters neurodevelopmental trajectories in mice: an integrated neuroimaging, behavioural, and transcriptional study

**DOI:** 10.1101/2020.12.03.406454

**Authors:** Elisa Guma, Pedro Bordignon, Gabriel A. Devenyi, Daniel Gallino, Chloe Anastassiadis, Vedrana Cvetkovska, Amadou Barry, Emily Snook, Jurgen Germann, Celia M.T. Greenwood, Bratislav Misic, Rosemary C. Bagot, M. Mallar Chakravarty

## Abstract

Prenatal maternal immune activation (MIA) is a risk factor for neurodevelopmental disorders. How gestational timing of MIA-exposure differentially impacts downstream development remains unclear. Here, we characterize neurodevelopmental trajectories of mice exposed to MIA induced by poly I:C either early (gestational day [GD]9) or late (GD17) in gestation using longitudinal structural magnetic resonance imaging from weaning to adulthood. Early MIA-exposure associated with accelerated brain volume increases in adolescence/early-adulthood that normalized in later adulthood, in regions including the striatum, hippocampus, and cingulate cortex. Similarly, alterations in anxiety, stereotypic, and sensorimotor gating behaviours observed in adolescence normalized in adulthood. In contrast, MIA-exposure in late gestation had less impact on anatomical and behavioural profiles. Using a multivariate technique to relate imaging and behavioural variables for the time of greatest alteration, i.e. adolescence/early adulthood, we demonstrate that variation in anxiety, social, and sensorimotor gating associates significantly with volume of regions including the dorsal and ventral hippocampus, and anterior cingulate cortex. Using RNA sequencing to explore the molecular underpinnings of region-specific alterations in early MIA-exposed mice in adolescence, we observed the most transcriptional changes in the dorsal hippocampus, with regulated genes enriched for fibroblast growth factor regulation, autistic behaviours, inflammatory pathways, and microRNA regulation. This indicates that MIA in early gestation perturbs brain development mechanisms implicated in neurodevelopmental disorders. Our findings demonstrate the inherent strength of an integrated hypothesis- and data-driven approach in linking brain-behavioural alterations to the transcriptome to understand how MIA confers risk for major mental illness.

## 1. Introduction

Prenatal brain development is a complex process orchestrated by interacting genetic, environmental, and immune factors. During this period offspring are highly vulnerable to a variety of risk factors for neurodevelopmental disorders that may only emerge later in childhood or adolescence (1–3). Epidemiological and preclinical evidence supports maternal infection as a risk factor for neurodevelopmental disorders such as autism spectrum disorder (ASD) and schizophrenia acting as a disease primer (1,4–6).

The effects of maternal infection are most often attributed to the maternal immune activation (MIA) rather than the specific pathogen. The increase in maternal proinflammatory cytokines disrupts the delicate immune balance between maternal and fetal environments, altering developmental processes (7–9). Studying the impact of MIA-exposure in animal model offspring has been critical to establishing causality between MIA during pregnancy and downstream neurodevelopmental disruptions (6,10,11). For example, MIA in pregnancy induces enduring behavioural, neuroanatomical, and transcriptional alterations relevant to schizophrenia and ASD (11–16).

However, neurodevelopmental processes and maternal cytokine responsiveness vary across gestation. Thus, the gestational timing of MIA-exposure may influence the nature and severity of disruptions in offspring (13,17–19). Epidemiological studies suggest exposure in early gestation confers the greatest risk for offspring (20–23). Nonetheless, there is significant variation in the literature with respect to the effects of MIA-timing. Previous animal studies comparing MIA-exposure in early (gestational day [GD] 9, ~the end of the human first trimester) or late (GD 17, ~the end of the human second trimester) report diverging neuroanatomical and behavioural phenotypes in adult offspring (17,24–26).

The emergence of neuroanatomical and behavioural abnormalities in the context of neurodevelopmental disorders is a dynamic process best characterized longitudinally. This is a strategy that has long been championed as a means for examining how variation from a “normative” path may lead to disordered brain development, potentially giving rise to neuropsychiatric complications (16,27–31). The effects of MIA are complex and may differ between regions, so whole brain strategies prioritizing developmental trajectories are particularly relevant in furthering our understanding of MIA-impact and moving beyond the many previous studies that have focused on isolated brain regions or behaviours in adulthood (16,25). Furthermore, given the current COVID-19 global crisis, gaining an understanding of the long term effects of MIA is critical, as the number of MIA-exposed offspring is expected to rise (32–34).

Here, we examine the impact of early (GD9) or late (GD17) prenatal MIA-exposure on developmental trajectories in mice using longitudinal whole-brain magnetic resonance imaging (MRI) and multi-behavioural characterization. We identify a window of deviation from normative trajectories, specifically the adolescent/early-adult period, and apply an integrative analysis at this timepoint to derive a pattern of linked brain-behaviour covariation to integrate anatomical changes in affected regions, including the hippocampus, anterior cingulate cortex, striatum, septal nucleus with affected behaviours, including anxiety, stereotypy, and sensorimotor gating. Greater behavioural impairments were associated with volumetric decreases in regions including the hippocampus, thalamus, and cerebellum, and larger volume in the cingulate cortex and striatum. We use transcriptional profiling to probe the molecular underpinnings of this emergent brain-behaviour pattern and find dysregulation of genes regulating fibroblast growth factor signaling, immune signaling and autistic behaviours in one of the most affected regions, the dorsal hippocampus. Using multiple dimensions of brain anatomy, behaviour and gene expression we construct a comprehensive understanding of how MIA-exposure, particularly in early gestation, shapes offspring neurodevelopment. Taken together, these findings provide key insights into how exposure to this risk factor in gestation may increase susceptibility for neurodevelopmental disorders later in the lifespan.

## 2. Results

### 2.1. Early and late gestational MIA-exposure differentially alter neurodevelopmental trajectory

To test the effect of MIA-exposure at GD9 or 17 on brain development across the human equivalent of childhood (postnatal day [PND]21), adolescence (PND38), early adulthood (PND 60), and adulthood (PND90), we collected longitudinal *in vivo* structural magnetic resonance images (MRI; 100μm^3^) (**Figure 1; section 4.2**). We examined developmental trajectories using linear mixed-effects models at each voxel in the brain, testing for group by age (cubic natural spline fit) interaction, with sex as a covariate, and random effects for litter and mouse **(section 4.2.3)**. This is akin to examining deviations from “normative” trajectories of brain development commonly performed in human neuroimaging studies (27,35). We observed the most significant deviations in developmental trajectories in early exposed poly I:C (POL E) offspring relative to saline (SAL) controls (for a quadratic age fit t=3.835, <1% false discovery rate [FDR]); POL E offspring had smaller brain volumes at P21, which then overshot between PND38 and PND60, and normalized at PND90. Many of these regions are implicated in neuro-psychiatric and -developmental disorders such as the hippocampus, subiculum, cingulate cortex, striatum, nucleus accumbens, and septal nucleus (36–40) **(Figure 2)**. Other important regions also affected include the periaqueductal gray and later developing regions such as the cerebellar vermis/crus I further detailed in **supplementary figure 4**. Offspring exposed to late poly I:C (POL L) had a flatter developmental growth trajectory relative to SAL (for a cubic age fit t=5.286, <1%FDR), including the nucleus accumbens, auditory cortex, reticular nucleus, subiculum and hypothalamus, whereas the amygdala volume decreased in later adulthood (**Figure 2 & supplementary figure 4**). These regions implicated in neuropsychiatric disorder show a somewhat different curve than that observed in the POL E group. POL E trajectories were significantly different from POL L, confirming that early MIA exposure had the largest effect on brain anatomy (**section 4.2.3; supplementary materials 2.5 & supplementary figures 9 & 10).** We also observed a significant monotonic increase in volume of cortical regions in POL E offspring (vs SAL) with a linear age fit; cubic age for POL E (vs SAL) and linear and quadratic age fits for POL L (vs SAL) were also significant, described in **supplementary materials 2.3 & 2.4 & supplementary figures 5-8**. We also explored sex differences as a post-hoc analysis, as our model comparison did not suggest it was the optimal model for our data; POL E (vs POL L) males were more affected than females in a number of regions, but none were observed with SAL as the reference group (**supplementary materials 2.6 & supplementary figure 11**).

**Figure 1.**
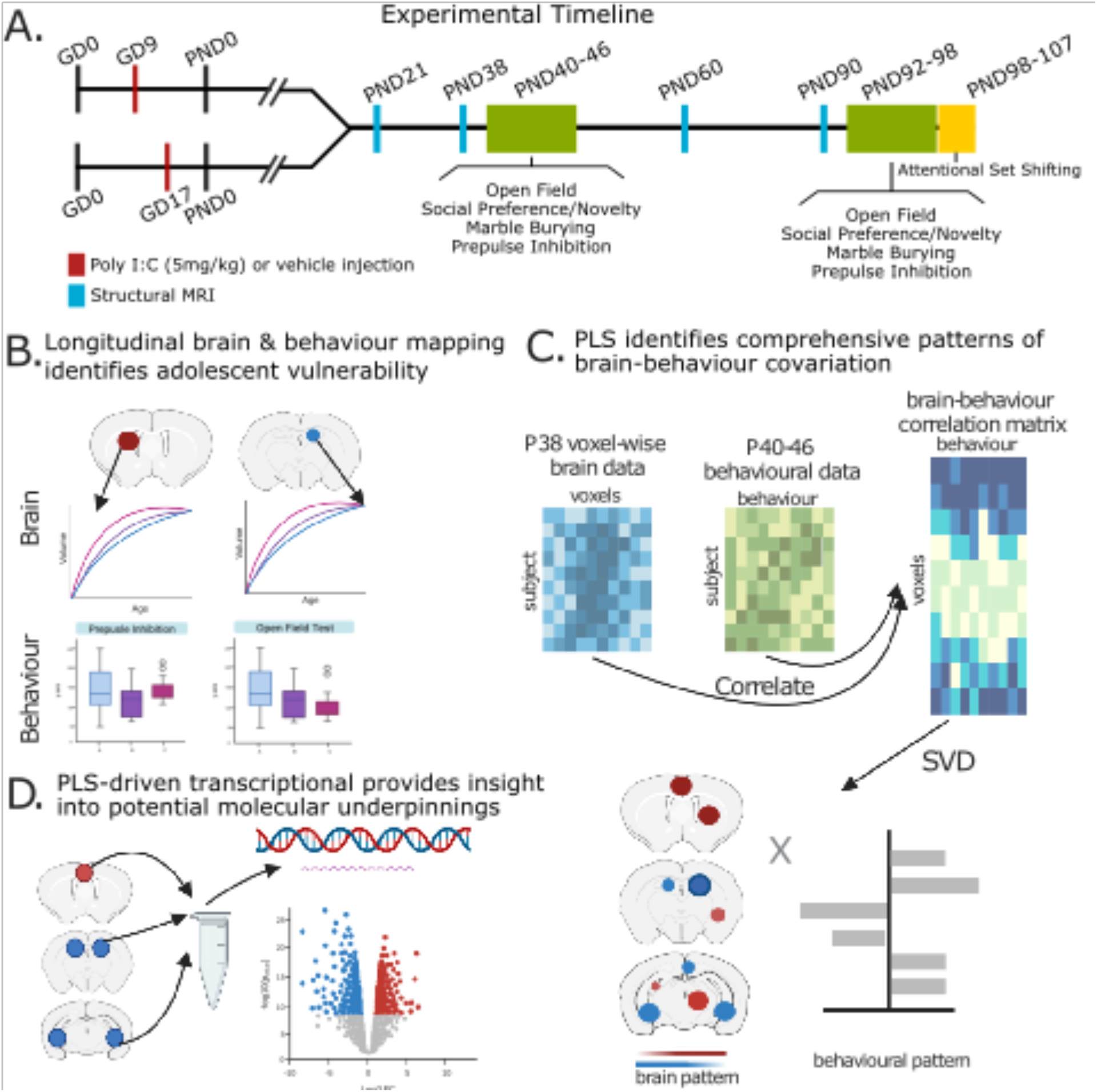
Experimental timeline. **A.** Pregnant dams were either injected (i.p.) with poly I:C (5m/kg) or vehicle (0.9% sterile NaCl solution) at gestational day (GD) 9 or 17 (red bars). Offspring were weaned and sexed on postnatal day (PND) 21. Longitudinal structural magnetic resonance imaging (MRI) was performed at PND 21, 38, 60, 90 (light blue bars). Two days following the PND38 (adolescence) and PND 90 (adulthood) scans, mice were assessed in open field test, social preference/novelty test (3 chambered social approach), marble burying task, and prepulse inhibition (green bars). The attentional set shifting task was also performed following the last adult behaviour (yellow bar). **B**. Univariate analyses were performed to assess group differences in neuroanatomy over time and behaviour for offspring exposed to MIA at GD 9, the early poly I:C (POL E) group, and offspring exposed at GD17, the late poly I:C group (POL L) group, relative to our combined vehicle exposed group, SAL (GD9 + 17). **C.** Partial least squares (PLS) was used to identify patterns of brain-behaviour covariation at the adolescent timepoint (PND38) where we observed our greatest group differences. This was used to identify regions of interest for RNA sequencing (**D**) used to probe potential molecular underpinnings of the observed changes. Figure made with BioRender https://biorender.com/.

**Figure 2.**
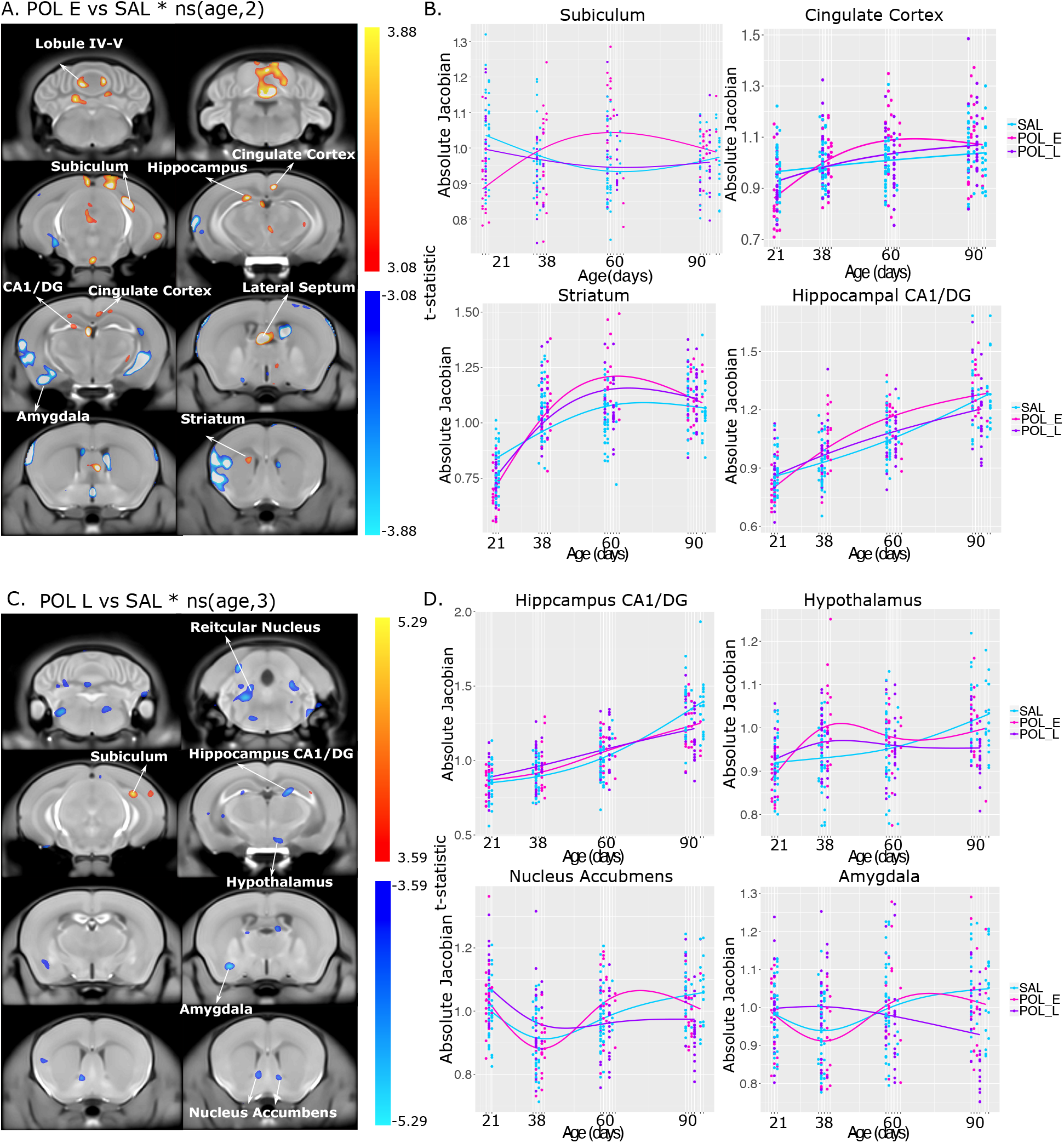
Developmental trajectories differ between early poly I:C group (POL E) vs saline controls (SAL) & the late poly I:C group (POL L) vs SAL (thresholded at 5% False discovery rate (FDR)) (**A**) t-statistic map of group (POL E vs SAL) by age (quadratic natural spline) thresholded between 5% FDR (bottom, t=3.08) and 1% FDR (top, t=3.83) overlaid on the population (second level) average (**B**) Plot of peak voxels (voxel within a region of volume change showing largest effect) selected from regions of interest highlighted (**A**), wherein age is plotted on the x-axis, and the absolute Jacobian determinants plotted on the y-axis. Here a value of 1 means the voxel is no different than the average, anything above 1 is relatively larger, and below 1 is relatively smaller. Ranges are not normalized to enhance comparison at each specific location in space. Trajectories are modeled as quadratic natural splines to reflect statistical modeling. (**C**) t-statistic map of group (POL L vs SAL) by age (cubic natural spline) thresholded between 5% FDR (bottom, t=3.59) and 1% FDR (top, t=5.29). (**D**) Plots of peak voxels as described in (**B**) with curves modeled as cubic natural splines to reflect statistics.

### 2.2. Early MIA-exposure induces behavioural alterations in adolescence

To capture the developmental emergence of behavioural abnormalities, we performed a battery of tests to assay schizophrenia- and ASD-relevant behaviours in the same animals that underwent neuroimaging (**section 2.1)** after the adolescent (PND38) and adult (PND90) scans. We assessed exploratory behaviour and anxiety (open field test), social behaviour (three chambered social preference task), stereotypy (marble burying task), and sensorimotor gating (prepulse inhibition to acoustic startle) (**Figure 1; section 4.3**).

In adolescence, POL E offspring traveled less than SAL in the anxiogenic center zone of the open field relative to total distance of the arena, however this effect did not survive multiple comparisons correction (uncorrected p values reported, and Bonferroni q values, ɑ= 0.05/11 = 0.0045; t=-2.294, uncorrected p=0.039, q=0.429; **section 4.3.2**; **Figure 3A**). They buried more marbles than SAL in the marble burying task, potentially suggesting greater stereotypy/anxiety (t=2.937, p=0.003, q=0.033; **Figure 3B**). Relative to SAL, POL E displayed a striking impairment in sensorimotor gating (t=-4.202, p=4.0 × 10^−7^, q=4.0 × 10 ^−6^; **Figure 3C**) across prepulse tones (i.e. no interaction between group and prepulse level: t=-0.995, p=0.321; **Figure 3D**). Surprisingly, adolescent offspring showed no impairments in social preference or novelty **Figure 3E**). Moreover, behavioural alterations were no longer present in POL E offspring in adulthood apart from a subthreshold impairment in social novelty behaviour (t=-2.369, p=0.0311, q=0.341; **Figure 3F**), wherein POL E mice spent more time exploring the familiar than the novel social target.

**Figure 3.**
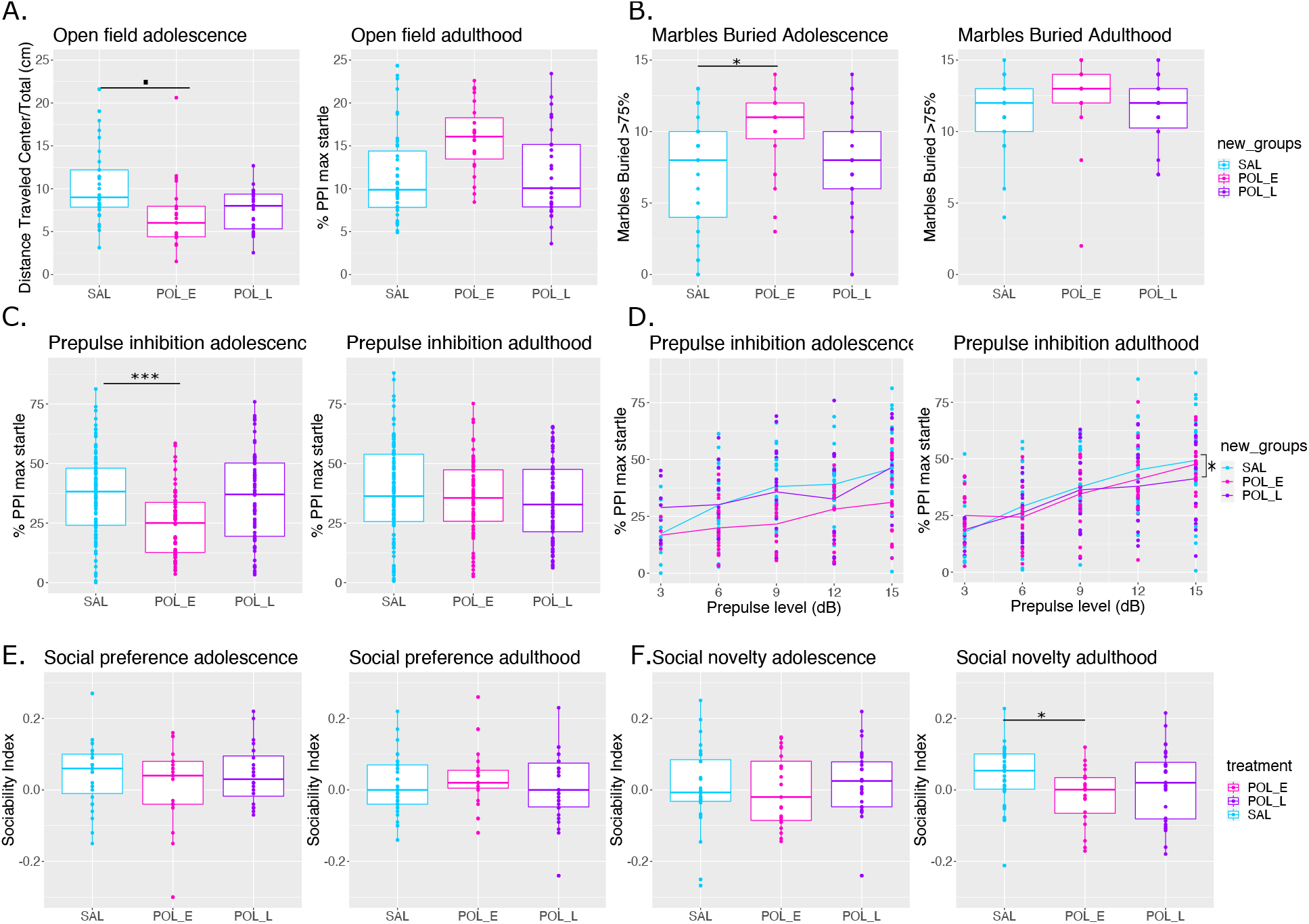
Early MIA-exposure induces transient behavioural impairments whereas late MIA-exposure does not affect behaviour. Behavioural results for adolescent (left) and adult (right) offspring from the three treatment groups: SAL (cyan), POL E (magenta), POL L (purple). For all boxplots the midline represents the median of the data, the box represents the interquartile range, with whiskers denoting the full range of the data. **(A)** In adolescence (left), POL E offspring travel less in the center zone (relative to the total distance traveled) at a subthreshold level (t=-2.294, p=0.039 not significant following Bonferroni correction). No statistically significant differences observed in adulthood (right). (**B**) Significantly more marbles are buried by POL E adolescent offspring (left; t=2.937, p=0.003) than SAL offspring. No group differences were observed in adulthood (right). (**C**) POL E offspring show a significant decrease in % prepulse inhibition based on maximum amplitude of startle reaction to startle tone (left; t=-4.202, 4.0 × 10 –7). This deficit is no longer present in adulthood (right). (**D**) %PPI is plotted for each group with increasing prepulse tone on the x-axis and %PPI based on maximum amplitude of startle reaction on the y-axis for adolescence (left) and adulthood (right). No significant differences in slopes are observed between groups, however the POL E offspring are impaired at all levels in adolescence. (**E**) No significant differences in sociability index for the social preference task (i.e. preference for novel mouse over nonsocial object) between groups at either adolescence (left) or adulthood (right). (**F**) No significant differences in sociability index for social novelty (i.e. preference for novel mouse over familiar mouse) between groups. A subthreshold impairment was observed in POL E adult offspring (t=-2.369, p=0.031). **ॱ** p<0.05; *p<0.0045 (Bonferroni correction threshold); ***p<0.0001

Adolescent POL L offspring did not show any significant behavioural alterations. Some subtle behavioural alterations emerged in adulthood, however they did not survive Bonferroni correction. Details in **supplementary section 2.7** and results summary in **supplementary table 3**. Only subthreshold effects were observed in the attentional set shifting task (ASST) performed only in adulthood, (**supplementary materials 2.7.1**& **supplementary figure 12)**. Post-hoc analyses of sex-differences are reported in **supplementary materials 2.7.2 & supplementary figure 13**.

### 2.3. Multivariate analysis of brain-behaviour data links variation in autism- and schizophrenia-related behaviours to volume changes in key brain regions

Based on the analyses described in **sections 2.1 & 2.2** we identified the adolescent/early adult period as one of greatest deviation from “normative” trajectories (particularly for the POL E group). Focusing on single modalities (neuroimaging or behaviour only) may obscure more complex relationships between brain and behaviour. To probe these potentially critical associations, we employed a data-driven technique, partial least squares (PLS), to perform multivariate mapping between whole-brain anatomical alterations and adolescent behavioural metrics across 4 different tests (PND 38; **Figure 1; section 4.4**). This analysis reveals patterns of covariation that link patterns of disordered brain development with disordered behaviours by identifying ‘latent variables’.

We identified two significant latent variables (LVs); LV1 described a pattern of brain-behaviour covariation (29% covariance explained, p=0.034), whereas LV2 described a brain pattern associated with sex and litter size (19% covariance explained, p=0.002) (**Figure 4A**; **supplementary materials 2.8 & supplementary figure 14)**. For this reason, we chose to focus on LV1. We observed a pattern of attenuated behavioural impairment, i.e. decreased locomotion and anxiety (open field test), more social interactions (social preference and novelty), and less impairment in sensorimotor gating (PPI) that was associated with larger volume of the ACC, somatomotor cortex, and striatum. A pattern of greater behavioural impairment, including increased anxiety and locomotion, fewer social interactions, and impaired sensorimotor gating was associated with smaller volume in the dorsal and ventral hippocampus, thalamic nuclei, and cerebellum (vermis, crus I and II) **(Figure 4 B&C**).

**Figure 4.**
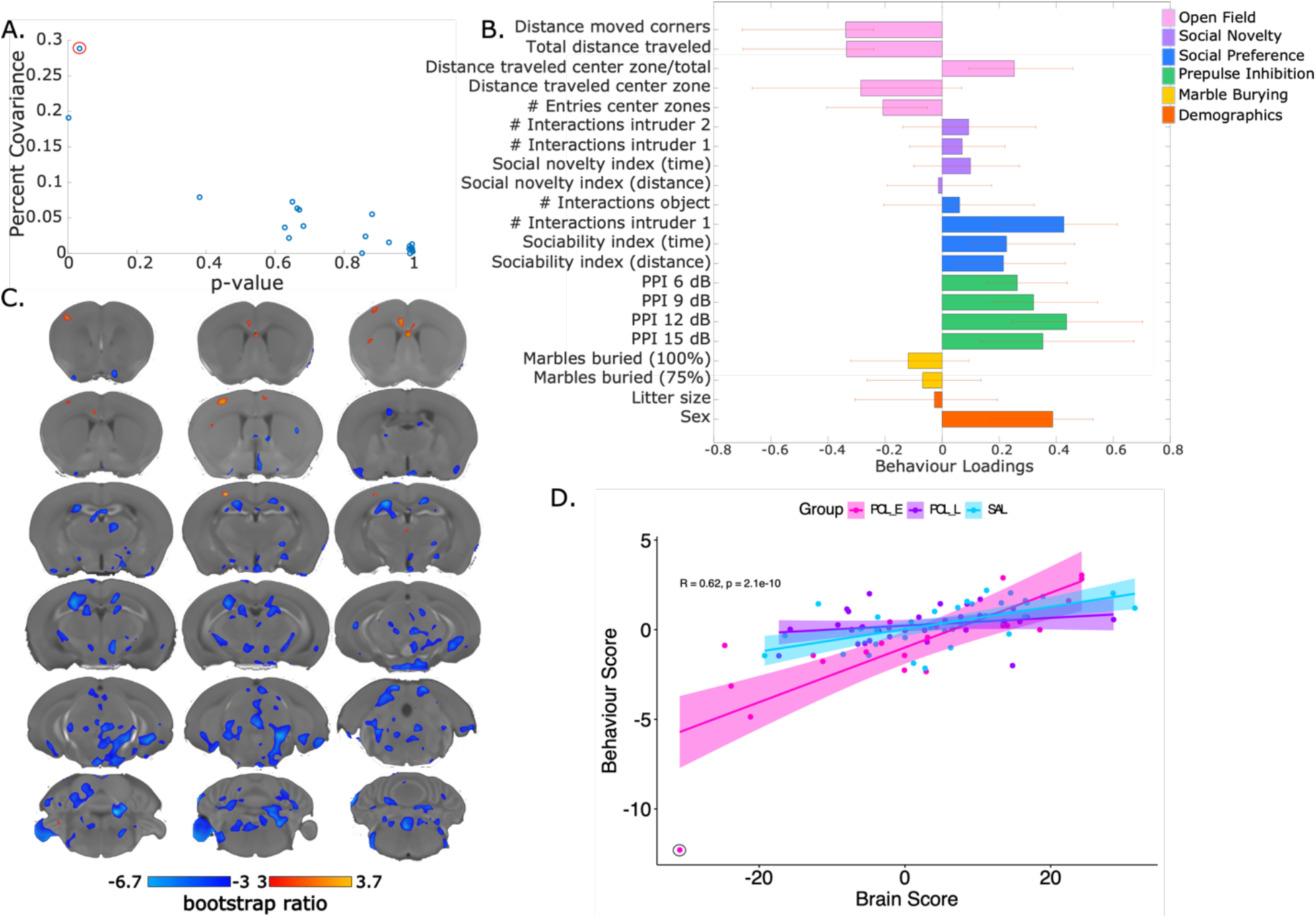
Partial least squares (PLS) analysis results for first latent variable (LV1). (**A**) Covariance explained (y-axis) and permutation p-values (x-axis) for all 21 LVs in the PLS analysis. LV1 is circled in red (p=0.034, %covariance=29%) and was chosen for subsequent investigation based on the covariance explained and behavioural relevance of results. (**B**) Behaviour weight for each behavioural measure included in the analysis showing how much they contribute to the pattern of LV1. Singular value decomposition estimates the size of the bars whereas confidence intervals are estimated by bootstrapping. Bars with error bars that cross the 0 line should not be considered. (**C**) Brain loading bootstrap ratios for the LV1 deformation pattern overlaid on the population average, with positive bootstrap ratios in orange-yellow (indicative or larger volume), and negative in blue (indicative of smaller volume). Colored voxels make significant contributions to LV1. **(D**) Correlation of individual mouse brain and behaviour score, color coded by treatment group with a trend line per group. Outlier on the behaviour score circled in dark gray. Early poly I:C (POL E) offspring (magenta) express this pattern more strongly than the saline controls (SAL) and late poly I:C (POL L) groups.

Inspecting correlations between the brain-behaviour weights for each mouse, colored by group, suggests that the POL E offspring load more strongly on this pattern than the other two groups, albeit not for all subjects (**Figure 4D; supplementary figure 15** shows the brain-behaviour correlation plot after removal of an outlier on behavioural loadings circled on plot). Finally, to determine how this pattern changes in adulthood, we applied brain-behaviour weights for LV1 computed in adolescence to the brain and behaviour data collected in adulthood. We observed a shift along the brain axis but not the behaviour axis, suggesting that, as mice age, changes in brain patterns are disconnected from changes in behavioural patterns (**supplementary figure 16**).

### 2.4. Early MIA-exposure induces transcriptional changes in adolescence

Based on our data-informed multivariate maps of brain-behaviour covariation, as well as *a priori* knowledge of regions implicated in schizophrenia- and ASD-related pathology (38,41–47), we identified regions in which to probe underlying transcriptional alterations, namely the ACC, dHIP, and vHIP. We first profiled patterns of differential gene expression in adolescent POL E (vs SAL E) mice (**Figure 1; section 4.5.2)**. Pooling all ROIs, we identified 962 genes significantly (q < 0.05) down-regulated and 668 genes upregulated in POL E relative to SAL mice. We observed many differentially expressed genes (DEGs; p<0.05) in the dHIP (246 down- and 131 upregulated, q < 0.05), with more subtle changes in the vHIP (37 down-, 12 upregulated, q < 0.05) and ACC (17 down-, 0 upregulated, q < 0.05) (**Figure 5A-C**). Several genes were significantly downregulated across all three ROIs, including *Nfkbia*, a key driver of the pro-inflammatory immune response for increasing cytokines production (48), as well as *Klf2*, *Ddit4*, and *Per1* (**Figure 5D & E)**. Full gene lists are available in **supplementary table 4.**

**Figure 5.**
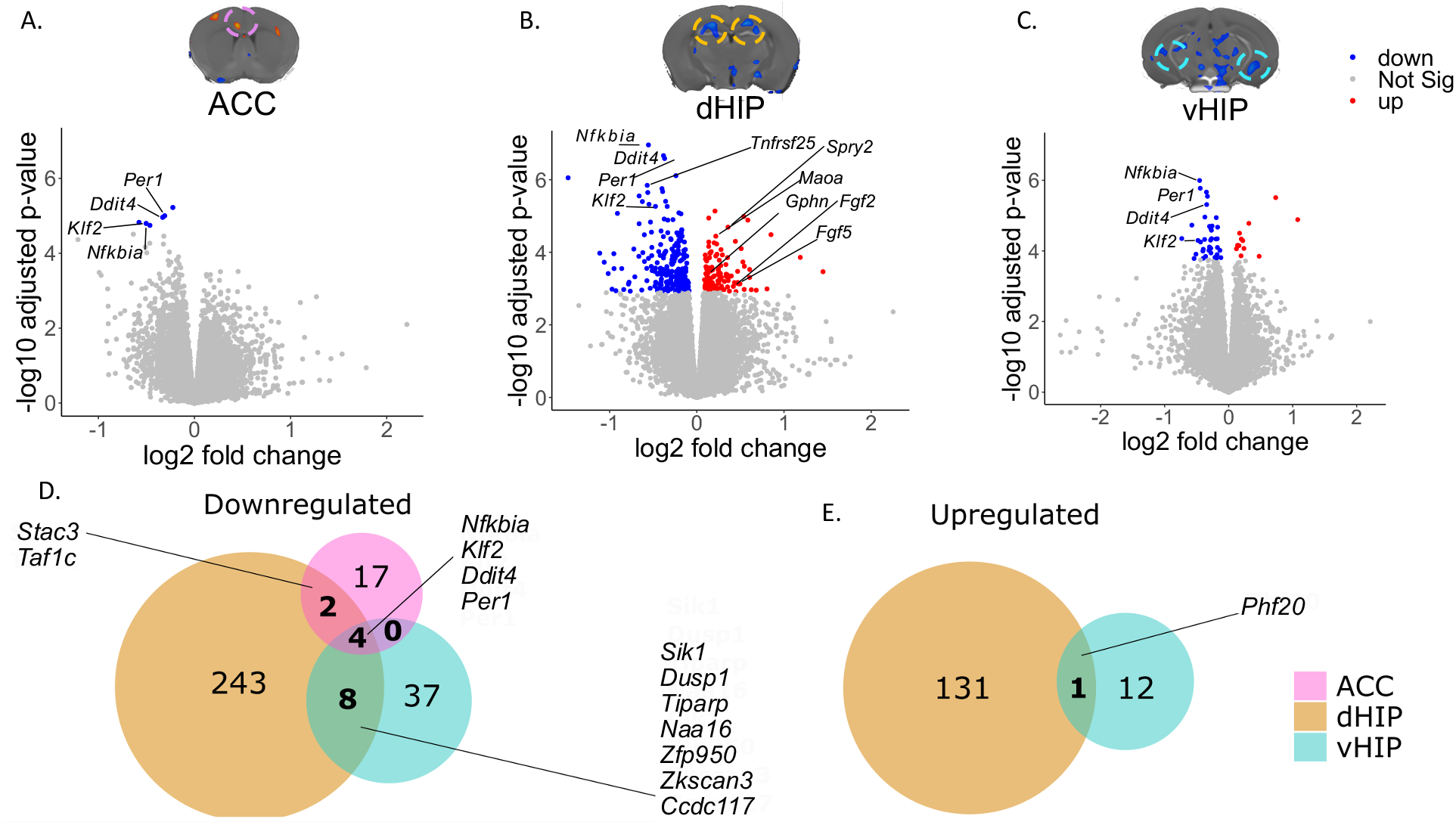
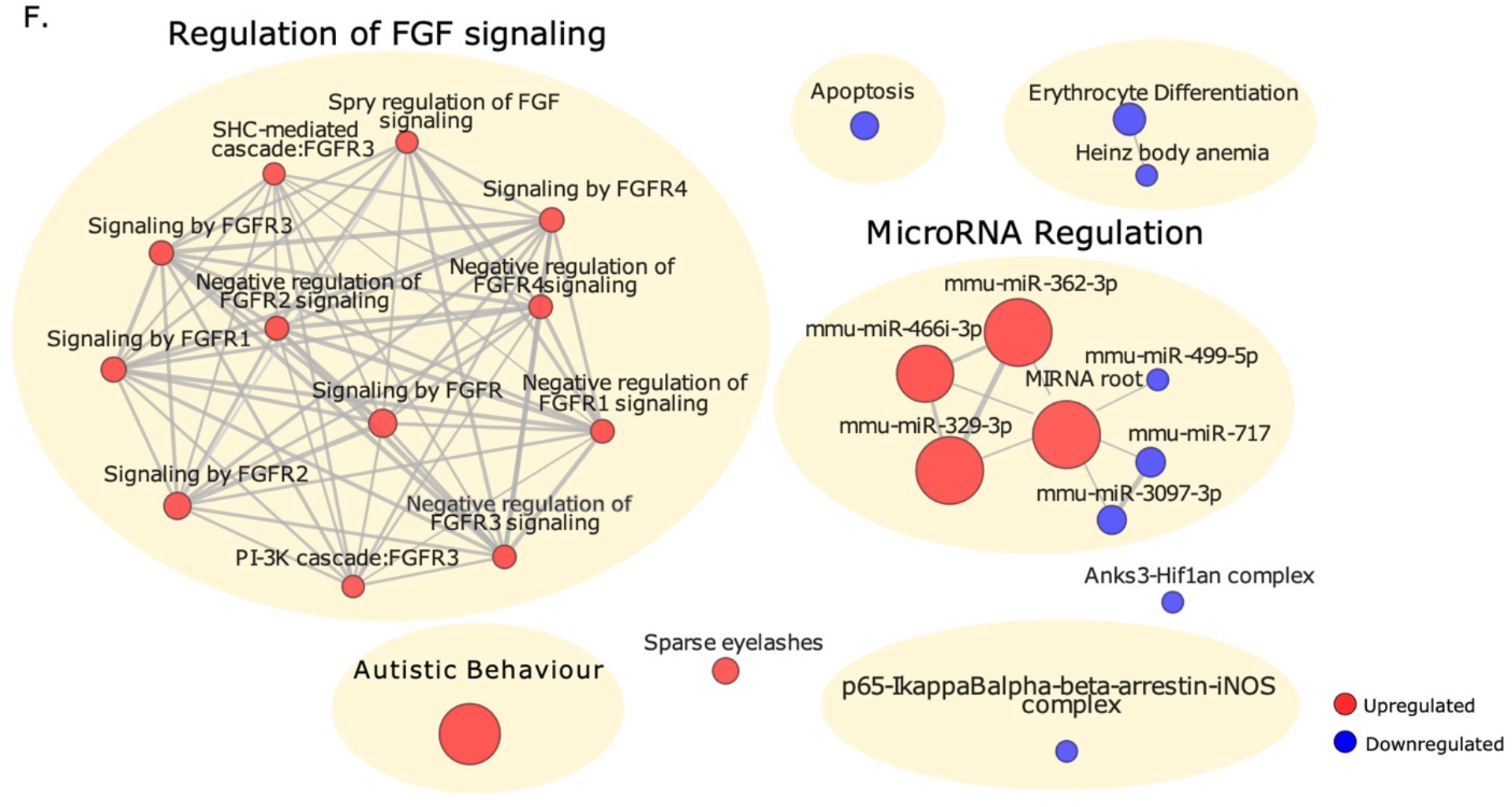
Transcriptional alteration in the adolescent brain at PND38 following MIA-exposure at GD9. Volcano plots for the ACC (**A**), dHIP (**B**), and vHIP (**C**) with significantly downregulated genes in blue, and upregulated genes in red. Genes that are either down- or upregulated in multiple ROIs (as shown in **D & E**) are highlighted in the volcano plots (on the left side). For the dHIP volcano plot (**B**) gene names on the right hand side are enriched for FGF signaling, and also identified in human postmortem hippocampal samples from individuals with schizophrenia. Venn diagram showing overlap in downregulated (**D**) and upregulated (**E**) genes per ROI. **F**. Gene enrichment analysis results for the dHIP, with upregulated enrichment in red and downregulated in blue. Upregulated genes were significantly enriched for FGF signaling, as well as autistic behaviours, sparse eyelashes, and microRNA regulation. MicroRNA regulation was also enriched for downregulated genes, as were apoptosis, erythrocyte differentiation, heinz body anemia, and p65-IKβKɑ-β-arrestin-iNOS-complex.

Pathway enrichment analysis of genes upregulated in dHIP **(section 4.5.3)** identified enrichment of fibroblast growth factor signaling (including FGF1, 2 and 3), an evolutionarily controlled signaling pathway critically involved in embryogenesis and synaptogenesis (49). Autistic behaviour and regulation of microRNAs (including miR-466i-3p, miR-362-3p, miR-329-3p) were also enriched. Downregulated dHIP genes were enriched for apoptosis, a critical neurodevelopmental process previously shown to be disrupted by MIA-exposure (50,51) and microRNAs (including miR-3097-3p, miR-499-5p, miR-717). Enrichment of NF-kappa-B inhibitor alpha signaling and erythrocyte differentiation enrichment, both implicated in the immune system, was also observed (52,53) **(Figure 5E; supplementary table 5)**.

In the vHIP, downregulated genes were enriched for IL-17 signaling pathway, previously shown to be critical for promoting an ASD-like phenotype in MIA-exposed mice (54), apoptosis, NF-kappa-B inhibitor alpha signaling, chronic myeloid leukemia and small cell lung cancer.

ACC upregulated DEGs were enriched for white fat cell differentiation and IL-1 signaling, also involved in pro-inflammatory signaling(55), based on wikipathways (https://www.wikipathways.org/index.php/WikiPathways). We investigated sex differences but no striking differences were observed (**supplementary materials 2.9.1**).

We compared identified DEGs to those previously identified by transcriptional profiling of post-mortem human ASD (prefrontal and temporal cortex (56)) and schizophrenia samples (pan-cortical (56), and prefrontal cortex and hippocampus (57)). Although disease gene lists were not significantly enriched in the mice, *FGF2* (significantly enriched in upregulated dHIP genes), *Tbx4*, and *Ccdc92* were upregulated in both the dHIP of our mice, and the hippocampus of individuals with schizophrenia (57). Further, there was some overlap in downregulated genes across all 3 ROIs and the pan-cortical SCZ sample, including *Nfrkb* (related to *Nfkbia* which was downregulated in all 3 ROIs), amongst others listed in **supplementary table 6**. This further supports the relevance of our findings to neuropsychiatric disorders.

To explore the transcriptional synchrony between brain regions following prenatal MIA exposure, we used RRHO to identify genome-wide (threshold-free) patterns and significance of overlap between gene expression profiles across pairs of brain regions. We observed the strongest overlap in genes downregulated in both the dHIP and vHIP (4932) **(supplementary figure 17A**), however, there was a degree of overlap between all pairwise comparisons. Coordinately upregulated genes across brain regions were enriched for myelin associated processes, oxidative phosphorylation, and mitochondrial function. Genes coordinately downregulated across all ROIs were enriched for RNA processing and transcriptional regulation. We report sex differences in gene overlap in **supplementary materials 2.9.2**& **supplementary figures 17B**.

## 3 Discussion

In this study, we demonstrate that integrating brain and behavioural phenotypes can uncover key relationships between aberrant development due to MIA-exposure and the putative transcriptional underpinnings. Specifically, we present whole-brain longitudinal neuroimaging and multi-behavioural phenotyping of mice prenatally exposed to MIA at two different gestational timepoints to evaluate developmental trajectories from childhood to adulthood, finding that early MIA induces the most robust changes. Integrating data from the developmental window of greatest vulnerability to characterize patterns of brain-behaviour covariation by multivariate analysis, we identified key brain regions and investigated transcriptional alterations in adolescence in the most affected group, POL E, revealing putative molecular underpinnings of the observed MIA-induced brain-behaviour patterns. Based on these analyses, we conclude that prenatal MIA-exposure early in gestation opens a window of vulnerability in adolescence/early adulthood characterized by an accelerated increase in brain volume and emergence of sensorimotor gating and stereotypy/anxiety-like deficits, both of which normalize in adulthood. These deviations may be driven by transcriptional changes in genes involved in FGF signaling, autistic behaviours, inflammatory pathways, and microRNA regulation particularly in the adolescent dHIP.

### 3.1. The case for longitudinal investigation

An accurate assessment of normal and abnormal brain development can be best achieved with a longitudinal approach. Our understanding of the emergence of neuropsychiatric disorders such as schizophrenia or ASD has been enhanced by longitudinal neuroimaging studies showing that deviations in brain development begin prior to disease onset (58,59). Additionally, longitudinal studies are more sensitive, requiring fewer participants than cross-sectional studies to detect subtle differences in brain structure (60) and better account for interindividual differences (61). Volumetric changes ascertained with MRI provide a robust endophenotype in animal models that facilitate comparison to human illness (62,63). Previous animal imaging studies from our own group have demonstrated the utility of a longitudinal design in the context of ageing, genotype associations, and treatments (64–68), while others provide important insight into early brain development (69) or sex differences in the developing brain (28).

Using our longitudinal approach we identified transient deviations in development due to MIA-exposure in early gestation which may have been missed had we focused solely on adulthood. This may, in part, explain why some cross-sectional studies examining MIA-induced neuroanatomical alterations have found differing results depending on age of testing (13). Future work should examine trajectories starting earlier and extending later in the lifespan.

### 3.2. Early MIA-exposure is associated with greater deviations in neurodevelopmental trajectories

MIA-exposure affected several regions often implicated in ASD or schizophrenia including the striatum, hippocampus and subiculum (36–38,70), amygdala (36,71), periaqueductal gray (72–74) (amongst others such as the somatosensory cortices (75–77) and septal nuclei (78,79)). The shape of the developmental trajectories is also of significant interest. The overshoot in adolescence and early adulthood is reminiscent of the brain or cortical overgrowth phenomenon observed in ASD (39,47,80,81)(39). Conversely, the altered hippocampal and ACC morphology is in line with psychosis spectrum disorders (45,46). Previous MRI-based studies examining GD9 MIA-exposure have identified microstructural alterations in similar regions such as the ACC, hippocampus, lateral septum, and ventral striatum, albeit later in the lifespan (25,26).

POL E offspring exhibited impairments in behaviours relevant to both ASD and schizophrenia, namely sensorimotor gating, stereotypy/repetitive behaviours, and subthreshold deficits in anxiety-like behaviours, consistent with previous reports of greater suppression of exploratory behaviours and sensorimotor gating impairments (17). However, we observed transient impairments in adolescence that were resolved in adulthood, whereas others have observed emergence of these deficits in adulthood, not adolescence (16,17). This may be due to a number of factors including the dose of immunogen, route of administration, mouse strain, and more (82).

Exposure late in gestation led to subtler alterations in brain morphology, with some regions such as the amygdala showing decreases in adulthood, not paired with behavioural impairments. Previous cross-sectional MRI studies examining the impact of GD17 MIA-exposure in adult offspring report white matter and cerebellar volume decreases, and enlarged 4th ventricles (25,83). Previous longitudinal studies examining MIA-exposure in mid-gestation (GD15) have identified enlarged lateral ventricles, often observed in MRI investigation of schizophrenia (36), as well as decreased cortical and hippocampal volume and altered microstructure all emerging in adulthood, providing further evidence for the impact MIA-timing on offspring outcomes (16,25,26,84). Behaviourally, memory and cognitive deficits have been observed in GD17 MIA-exposed adult offspring (17,24,85), whereas we only observed subthreshold deficits in cognitive performance in the ASST task. Importantly, previous work comparing the effects of GD9 and 17 MIA-exposure also observed diverging effects on offspring brain development (17,25,86). Taken together, these findings indicate increases in maternal cytokine levels due to MIA that occur early in gestation may have more profound effects on offspring development. However, they also support the theory that MIA is a disease primer, as exposure to this risk factor alone does not seem to be sufficient in inducing long-term deficits later in the lifespan (5,11,87,88).

Although speculative, it is worth considering what neurodevelopmental processes may be altered by the MIA-exposure at the gestational epochs we chose. At GD9 the developing brain is colonized by microglia, and neuronal and immune cell migration, neurogenesis, and cortical plate formation are being initiated (3,89,90). At GD17 the organization of cortical layers and the hippocampus are underway, as are synaptogenesis, gliogenesis, and apoptosis (1,5,91). It remains unclear to what extent MIA-exposure during these periods influences downstream neurodevelopmental changes across the differing scales of brain architecture. Previously, MIA exposure at GD9 has been associated with an increased density of activated microglia both in the embryo and adolescent brain; this may in turn interfere with synaptic pruning and circuit formation leading to aberrant neurodevelopment (92–94). Alterations to myelin-related structure and gene expression have been observed in GD17 MIA-exposed offspring (83). Finally, MIA in mid-gestation (GD12-15) has been associated with aberrant neurogenesis, reductions in the number of cortical neural stem cells, and reduced dendritic spine density (95,96). Future work examining MIA at different gestational times (i.e. mid-gestation) with the integrative approach we developed may further our understanding of how MIA disrupts brain development.

### 3.3. Identifying brain-behaviour associations

One major limitation when examining neurodevelopmental phenotypes in either human or animal models is the tendency to assign phenotypes at the level of a single structure, or to examine associations between these structures and a specific symptom or behaviour (97). Indeed, these strategies disregard the ever-increasingly acknowledged network-like architecture of the brain and the inherent relationships to behaviour (98,99). In contrast, multivariate strategies, such as PLS, move beyond simplistic associations to better understand brain-behaviour relationships (100–102). In the context of small animal imaging, PLS provides a novel, streamlined method to assess cross-sectional brain-behaviour patterns from large cohorts of deeply phenotyped mice. This analysis, seldom applied to preclinical studies, has been widely applied to human neuroimaging studies investigating, for example, the relationship between neuroanatomy and gene expression (103) or clinical and cognitive symptoms in schizophrenia (104). This integrative analysis permits inclusion of multiple measures from the same individual while accounting for their potential inter-relatedness. Further, it balances hypothesis driven experimental choices (brain regions, behaviours tested, etc) with hypothesis-free data-driven investigation of their relationships. Finally, it may promote cross-species translation by improving methodological homology between human and rodent neuroimaging studies.

### 3.4. Neuroimaging-driven RNA sequencing reveals potential molecular underpinnings of brain-behaviour relationships

We observed significant enrichment of gene ontology terms for FGF signaling pathways, as well as autistic behaviour, and microRNA regulation amongst upregulated dHIP genes. Interestingly, we also saw enrichment of immune pathways such as IL-17 in the vHIP, IL-1 in the ACC, and NFK-B in the d- and vHIP in line with previous findings (54). This provides further evidence for immune system dysregulation following MIA-exposure, which may interfere with neurodevelopmental processes such as neuronal migration, microglial function, and immune system development (8,105,106).

FGFs are signaling proteins that influence the development and repair of most mammalian tissues (107,108). They play a critical role in embryonic development, influencing the growth and patterning of several brain structures, cell survival and proliferation, neurogenesis and neuronal repair (109,110). In preclinical studies, FGF signaling inhibition has been observed to disrupt cortical gyrification, neural progenitor development in the subventricular zone of the developing cerebral cortex (111), hippocampal excitatory and inhibitory synaptogenesis and cell maturation (112,113), and left-right symmetry, all phenomena observed in schizophrenia and autism (114). Enhanced FGF signaling has been associated with beneficial effects, such as reduced anxiety, enhanced neurogenesis, and reverses hippocampal and cortical atrophy (115,116). Dysregulation of FGF signaling has been proposed to increase vulnerability to ASD, producing variation in brain growth and cortical circuit formation (117,118). Further, a few of the FGF signaling genes we observed to be regulated have also been implicated in schizophrenia in human post-mortem brains (57). FGF receptor 2 single-nucleotide polymorphisms associate with schizophrenia (108,119,120); this gene was significantly upregulated both in the dHIP of our early MIA-exposed mice and the hippocampi of individuals with schizophrenia (57).

### 3.5. Limitations

The results and discussion of this paper should be considered in light of their limitations. No animal model can fully recapitulate the human condition, therefore not all observations made here may apply to human pathology and brain development (89). However, we do observe interesting parallels between our MIA-exposed mice and psychiatric patients’ altered trajectories of brain development, behaviour, and transcription. Further, studies investigating the impacts of MIA exposure in humans also report that earlier exposure induces more serious downstream effects on offspring (see (13) for a review). Although both the neuroimaging and behavioural studies conducted were longitudinal, our multivariate and transcriptional analyses were cross-sectional. Future work should be done to extend these analyses across the lifespan. We investigated sex differences; however we did not detect statistically significant interactions in neuroanatomy or behaviour. Although we collected data for both male and female offspring, we may still have been underpowered to detect potentially subtle sex-specific effects. When comparing our two POL groups directly, we did observe greater alterations in male POL E offspring, which may indicate existence of subtle sex-effects (121,122). Surprisingly we did not observe any strong social deficits, often reported in MIA-offspring and central to ASD pathology (11,54,123). This may be a function of MIA-timing, age of investigation, dose of immunogen used, or route of administration. In contrast to our findings, previous studies observed MIA-offspring deficits in adulthood (16,17); it is possible that these may emerge later in the lifespan or following exposure to an additional risk factor.

### 3.6. Conclusions

We comprehensively examined the effects of prenatal MIA-exposure, a known risk factor for neuropsychiatric disorders, at two different gestational timepoints, on offspring brain and behaviour development using a robust translational measure: *in vivo* rodent imaging. We applied multivariate statistical analyses to integrate these modalities, leveraging this to investigate underlying transcriptional changes in the group and age at which we detected the greatest changes. Taken together, these findings suggest that prenatal MIA-exposure early in gestation may interfere with critical neurodevelopmental processes more so than exposure in late gestation. This leads to transient deviations in brain and behaviour development in adolescence and early adulthood, potentially increasing susceptibility to other risk factors, but, which, in the absence of subsequent challenge, normalize later in adulthood. These may be linked to altered transcription of genes involved in FGF signaling and inflammatory pathways.

## 4. Methods

### 4.1. Animals, Breeding & Maternal Immune activation protocol

Timed-mating procedures were used to generate pregnant dams, who were injected intraperitoneally with either poly I:C (POL; P1530-25MG polyinosinic– polycytidylic acid sodium salt TLR ligand tested; Sigma Aldrich) (5mg/kg) or vehicle (SAL; 0.9% sterile NaCl solution) at GD 9 or 17 resulting in 4 groups: POL E (5 dams), SAL E (4 dams), POL L (6 dams), SAL L (5 dams). Experimental design is outlined in **Figure 1**(see **Supplementary materials 1.1** for breeding, model, birth and weaning details). We confirmed the immunostimulatory potential of our poly I:C in a separate set of dams (details for procedure in **supplementary materials 1.6 and results in supplementary materials 2.1 & supplementary table 2**).

### 4.2. Magnetic resonance imaging

#### 4.2.1. Acquisition

Longitudinal T1-weighted (100 μm^3^) structural MRI scans were acquired *in vivo* at PND 21 (~childhood), 38 (~adolescence), 60 (~young adulthood), and 90 (~adulthood) (89,90) at the Douglas Institute in MIA- or SAL-exposed offspring (7 Tesla Bruker Biospec 70/30; matrix size of 180 × 160 × 90; 14.5 minutes, 2 averages) (64–66,68). MnCl2 (62.5 mg/kg) was used for contrast enhancement (60,124), and isoflurane to anesthetize the mice (5% induction, 1.5% maintenance) (**Supplementary materials 1.3** for more details).

#### 4.2.2. Image processing

Images (n=376) were exported as Digital Imaging and Communications in Medicine (DICOM), converted to the Medical Imaging NetCDF (MINC) format, preprocessed to enable downstream analyses, and visually inspected for quality control (QC; n=27 scans were excluded https://github.com/CoBrALab/documentation/wiki/Mouse-QC-Manual-(Structural)).

Preprocessed images were used as inputs for the two-level Pydpiper registration toolkit to perform longitudinal deformation based morphometry analysis (https://wiki.mouseimaging.ca/pages/viewpage.action?pageId=1868779) (125,126). Briefly, in the first level, affine and non-linear registration was used to create a subject-average per mouse (by registering all scans for one mouse). In the second level all subject averages were registered to create a study average and a common space for statistical analysis (125) (**supplementary figure 1** for workflow). Quality control (QC) was performed through visual inspection to ensure that registrations worked as expected. Final numbers per group/timepoint are presented in **table 1**

**Table 1.** Sample per timepoint following quality control. Postnatal day (PND); poly I:C (POL); saline (SAL); late (L; gestational day 17 exposure); early (E; gestational day 9 exposure); male (M); female (F).

|  | PND 21 scan | PND 38 scan | PND 60 scan | PND 90 scan |
| --- | --- | --- | --- | --- |
| **SAL E + L (9 litters)** | 37 (19M/18F) | 38 (20M/18F) | 41 (21M/20F) | 40 (20M/20F) |
| **POL E (5 litters)** | 21 (8M/13F) | 21 (9M/12F) | 20 (8M/12F) | 20 (8M/12F) |
| **POL L (6 litters)** | 27 (12M/15F) | 29 (13M/16F) | 27 (14M/13F) | 28 (14M/14F) |

Both relative (nonlinear deformations with residual affine removed) and absolute (linear plus nonlinear deformations) Jacobian determinants (127) of the deformation fields of the first level were resampled into the second level final average space in order to perform statistics, and blurred with a 0.2 mm full-width-at-half-maximum 3D Gaussian to better conform to Gaussian assumptions for downstream statistical testing (125,126). Further detail in **supplementary materials 1.3** and **supplementary figure 1**.

#### 4.2.3. Statistical analysis

Statistical analyses were performed using the R software package (R version 3.5.1, RMINC version 1.5.2.2 www.r-project.org). Linear mixed-effects models (lme4_1.1-21 package) were chosen for our longitudinal analyses as they are robust to missing data, and allow for fitting both fixed and random effects for the same subject over time. To select the most appropriate natural spline fit for our age term, we compared increasingly complex models with the log-likelihood ratio test at every voxel in the brain following similar methodology presented in previous studies (**Supplementary materials 1.7.1**) (28,69). Modeling age with a cubic natural spline fit best explained the variance in our data, with group and sex as fixed effects, and litter and mouse id as random intercept effects (**supplementary section 2.2** and **supplementary figure 3**). SAL E and L offspring were merged into a single group after ensuring they were not significantly different (**Supplementary materials 1.7.2**). A voxel-wise linear mixed-effects model was applied to the first level absolute Jacobian determinant (to examine within-subject trajectories) to assess the effect of MIA-exposure on development with an age (cubic natural spline fit) by group interaction, covarying for sex (fixed effects), while subject and litter were modeled as random intercept effects. The model was run again with POL L as the reference group in order to compare POL E and POL L directly. The False Discovery Rate (FDR) (128) correction for multiple comparison was applied to our statistical tests. As a follow up analysis, we investigated whether there was a sex by group by age (cubic nonlinear spline) interaction (**supplementary materials 2.3**).

### 4.3. Behavioural testing

#### 4.3.1. Behavioural protocols

Tests were performed following the adolescent (PND38) and adult (PND90) scans, with a 2 day resting phase between tests and 1 hour habituation to the behavioural room prior to the test. Videos were analyzed using the Ethovision XT 12 tracking system (Noldus Information Technology, Leesburg, VA, USA).

Briefly, the **open field test** was used to assess exploratory behaviour and anxiety by comparing the distance traveled in the anxiogenic center zone (40% of a 45×45 cm^2^ grey plastic arena) compared to the corners and edges. Mice were allowed to explore the arena for 15 minutes under bright light.

For the **three chambered social approach** task, mice were habituated (10 minutes) under red light to a three-chamber plastic box (26 (l) × 21.6 (w) × 21.6 (h) cm) with divider panels that have open doors, with a wire container (9.5 (h) 7.6 (d) cm) in the two extreme chambers. To measure social preference (10 minutes), interactions with an age-matched same-sex stranger mouse were compared to interactions with a non social object (each under wire containers); to measure social novelty, interactions were compared between the same stranger mouse and a novel stranger mouse (which replaced the non social object).

The **marble burying task** was used to assess stereotypic and repetitive digging behaviour. We measured how many of 15 equidistantly spaced marbles mice buried in 30 minutes under standard wood chip bedding (~7 cm deep layer) in a standard home cage (28 (h) × 17 (w) × 12.7 (h) cm); see (129).

Sensorimotor gating to acoustic startle was measured with the **prepulse inhibition task** using commercially available startle chambers (San Diego Instruments, San Diego, CA). Startle response to an acoustic stimulus (120 dB) was compared to trials in which the startle stimulus was preceded by a 30ms prepulse stimulus ranging from 3-15 dB above background noise (73-85 dB; 5 trials/stimulus, 3dB increments).

An additional cognitive flexibility and reversal learning measure (**attentional set shifting task [ASST]**(130)) was performed following PND90 behaviours to determine if there were lasting effects of the MIA-exposure, as previously reported in adult MIA-exposed offspring in late gestation (17) (**supplementary materials 1.4** for details). Mice were perfused following the last behaviour (**supplementary materials 1.5**).

#### 4.3.2. Statistical analysis

We used linear mixed-effects models for adolescent and adult behavioural data, with group and sex (SAL and male as references) as fixed effects, and litter as a random effect to account for possible litter variability. A Bonferroni correction was applied (5 tests in adolescence, 6 in adulthood: ɑ= 0.05/11 = 0.0045 set as significance threshold, uncorrected p-values, and corrected q-values reported). As with the neuroanatomy, a follow-up assessment of sex differences was carried out (**supplementary materials 2.7.2.**).

### 4.4. Partial Least Squares Analysis

Partial Least Squares (PLS) is a multivariate analysis which allows us to find the optimal weighted linear combination of two variables (voxel-wise DBM and behavioural metrics) that maximally covary together (131–133). A covariance matrix was computed from the z-scored brain (voxel-wise DBM measures) and behaviour (relevant behavioural metrics, PND38 tests, sex, and litter) matrices. Singular value decomposition was applied to the covariance matrix (134) to yield a set of orthogonal latent variables (LVs). This generates a set of ‘brain weights’ and ‘behaviour weights’ describing how each voxel or behavioural variable, respectively, weighs onto a given LV, and a singular value, describing the proportion of covariance explained by the LV. Brain and behaviour weights were projected onto individual subject data to generate subject-specific brain and behaviour scores. Permutation testing (n=1000 repetitions) was used to assess the statistical significance of each LV and bootstrap resampling (n=1000) was applied to assess the contribution of original brain and behaviour variables to each LV. Bootstrap ratios were thresholded at values corresponding to a 95% confidence interval (131,135). Further details can be found in **supplementary materials 1.7.3**.

### 4.5. Transcriptional analysis

#### 4.5.1. Brain Extraction and RNA Isolation

A separate cohort of mice (PND38; POL E n=6M/6F, 5 litters, SAL E n=6M/6F, 6 litters) were euthanized in their home cage, brains were rapidly extracted and placed in chilled PBS and sliced (1 mm thick sections); tissue was punched from the ACC (bregma+0.14 mm), dHIP (bregma -2.80 mm), and vHIP (bregma -3.08 mm) and flash frozen on dry ice (death to freezing all samples < 3 minutes). RNA extraction was performed using ReliaPrep Tissue RNA Miniprep system (Promega). RNA quantity and integrity was analyzed using Nanodrop (Thermo Fisher) and Bioanalyzer (Agilent). The samples were randomized during library preparation and sequencing was performed on Illumina NovaSeq 6000 at the McGill University Génome Québec Innovation Centre.

#### 4.5.2. Differential expression

Read counts per gene were preprocessed (**supplementary materials 1.8.1 & supplementary figure 2**). Differential expression (DE) between POL E and SAL E offspring was calculated using the limma-voom pipeline from Bioconductor’s limma package (Version 3.40.6) (136). A linear mixed model was used to account for the repeated measurements from each subject to assess DE between groups at each region of interest (ROI) (covarying for sex), as well as global differences between groups (covarying for sex and ROI), correcting for multiple comparisons with FDR (128). A follow up analysis was performed on each sex separately to assess potential sex differences (**supplementary materials 1.8.2**). Rank rank hypergeometric overlap (RRHO) was used to evaluate the similarity of genome-wide DE in pairs of regions by determining the degree of statistical enrichment using hypergeometric distribution while sliding across all possible thresholds through two ranked lists (137) (https://systems.crump.ucla.edu/rankrank/rankranksimple.php; see **supplementary materials 1.8.3).**

#### 4.5.3. Pathway Enrichment, Rank Rank Hypergeometric Overlap, and Disease Gene Analysis

Pathway enrichment analysis was performed using g:Profiler (138,139) to identify pathways with significantly enriched or overrepresented genes from our gene lists ranked by the-log10P value relative to a background gene list. Pathways include Gene Ontology (GO) terms (biological process, cellular components, molecular function), pathways (Reactome, KEGG), networks, regulatory motifs, and disease phenotypes (138). Enrichment significance was assessed using g:GOSt, applying the hypergeometric distribution. Multiple comparisons correction was performed using g:Profiler’s tailor-made algorithm g:SCS (140) (**supplementary materials 1.8.4**). To measure concordance of differential gene expression patterns between POL E and SAL E across ROIs and between sexes we applied a rank rank hypergeometric overlap test (RRHO) (Cahill et al., 2018); a pathway enrichment analysis was performed on overlap lists (**supplementary materials 1.8.3**).

Using the Bioconductor GeneOverlap package (https://www.bioconductor.org/packages/release/bioc/html/GeneOverlap.html) in R, dHIP and vHIP DEGs were compared to those identified by human post-mortem studies in the HIP of individuals with schizophrenia (57). Similarly, all our ROIs were compared to schizophrenia and ASD pancortical DEGs (56) **(supplementary materials 1.8.5**).

## Acknowledgements

We would like to thank the Ludmer Center of neuroinformatics and mental health (http://ludmercentre.ca/) at McGill University for their support with the transcriptional analysis. Additionally, we would like to thank Gülebru Ayranci, PhD for her help with MRI acquisition, and the Douglas Animal Facility staff for their support with animal care. Finally, we would like to thank Dr’s Bruno Giros and Salah El Mestikawy for lending us their centrifuge.

## Abbreviations

MIA: maternal immune activation
GD: gestational day
PND: postnatal day
POL: poly I:C
SAL: saline
E: early (GD9)
L: late (GD17)
OFT: open field test
PPI: prepulse inhibition
SOPT: social preference test
SONT: social novelty test
ASST: attentional set shifting task
FDR: false discovery rate
PLS: partial least squares
LV: latent variable
ROI: region of interest
ASD: autism spectrum disorder
FGF: fibroblast growth factor
DEG: differentially expressed gene
RRHO: rank rank hypergeometric overlap
QC: quality control
ACC: anterior cingulate cortex
dHIP: dorsal hippocampus
vHIP: ventral hippocampus

