## Supplementary Materials and Figures for "Early or late gestational exposure to maternal immune activation alters neurodevelopmental trajectories in mice: an integrated neuroimaging, behavioural, and transcriptional study"

### 1. Supplementary Methods

#### 1.1. Animals & Maternal Immune activation protocol

All procedures were approved by McGill University's Animal Care Committee under the guidelines of the Canadian Council on Animal Care. C57BL/6J mice were bred in our facility under a 12 hour light cycle (8am-8pm), with food and water access *ad libitum*. Females and males of breeding age (8-12 weeks) were placed in new cages (1:1 ratio) for up to 2 days until seminal plug is observed. This was considered gestational day (GD) 0. Each female was weighed and moved to a new cage. Animals were weighed again on injection day to confirm pregnancy.

C57BL/6J mice bred in our facility were used throughout the study. Pregnant dams were randomly assigned to one of four treatment groups: (1) poly I:C (P1530-25MG polyinosinic–polycytidylic acid sodium salt TLR ligand tested; Sigma Aldrich) (5 mg/kg, intraperitoneally) at gestational day (GD) 9 (n=5), (2) vehicle (0.9% sterile NaCl solution) at GD9 (n=4), (3) poly I:C at GD17 (n=6), or (4) vehicle at GD17 (n=5). GD9 corresponds roughly to the end of the first trimester in human gestation and GD17 corresponds to the end of the second trimester (Semple et al. 2013; Clancy et al. 2007; Meyer et al. 2006).

#### 1.2. Experimental design

Offspring were weaned and sexed at postnatal day (PND) 21 and housed 2-4 per cage. Up to three offspring of each sex were kept per litter. Offspring exposed to prenatal MIA via viral mimetic poly I:C or vehicle were examined using structural MRI (100um isotropic) at PND 21, 38, 60, and 90 at the Douglas Institute (7 Tesla small animal MRI, Bruker). Assessment of exploratory behaviour (open field test), social behaviour (social preference), stereotypic behaviour (marble burying task), and sensorimotor gating (prepulse inhibition) was performed following scans at P38 (adolescence). All behaviours were repeated after the P90 scan (adulthood) with the addition of a cognitive flexibility measure (attentional set shifting).

#### 1.3. Magnetic Resonance Imaging acquisition and processing

*Acquisition details:* Twenty-four hours prior to each MRI scan, mice were injected with $MnCl_2$ (62.5 mg/kg) for contrast enhancement. Anesthesia was induced with 5% isoflurane in oxygen and maintained with 1.5% isoflurane during the scan. Scans were conducted in a 7 Tesla Bruker, 30 cm bore magnet with AVANCE electronics. A 3D FLASH (Fast, Low Angle SHot) sequence was used with TE/TR of 4.5ms/20ms.

*Preprocessing details:* T1-weighted scans were preprocessed by stripping native coordinates, flipping left-right to maintain fidelity, denoising, correcting (Friedel et al. 2014; Avants et al. 2008) inhomogeneities in the bias field using the N4 algorithm (Tustison et al. 2010), and registering in LSQ6 alignment (i.e. 6 degrees of freedom are allowed for imagine alignment: translations and rotations along x, y, and z dimensions) (Ashburner and Friston 1998; Friedel et al. 2014; Collins et al. 1994). Visual quality control (QC) was performed to exclude any scans that had artifacts or signal dropoff that would prevent accurate registrations; subjects who had only 1 of 4 usable scans were also excluded from further processing (n=27 excluded; https://github.com/CoBrALab/documentation/wiki/Mouse-QC-Manual-(Structural)).

*Registration details:* Linear LSQ12 registration was applied to the preprocessed LSQ6 images using 12 degrees of freedom are allowed including those in LSQ6 as well as 3 scales and 3 shear parameters (Friedel et al. 2014; Lerch et al. 2008; Kovačević et al. 2004; Avants et al. 2008; Avants et al. 2011). Relative Jacobian determinants explicitly model only the non-linear part of the deformations and remove residual global linear transformations (attributable to differences in total brain size). Absolute Jacobians (without removal of overall linear transformations) were used to better determine what localized changes in volume are attributable to global changes in volume (Chung et al. 2001).

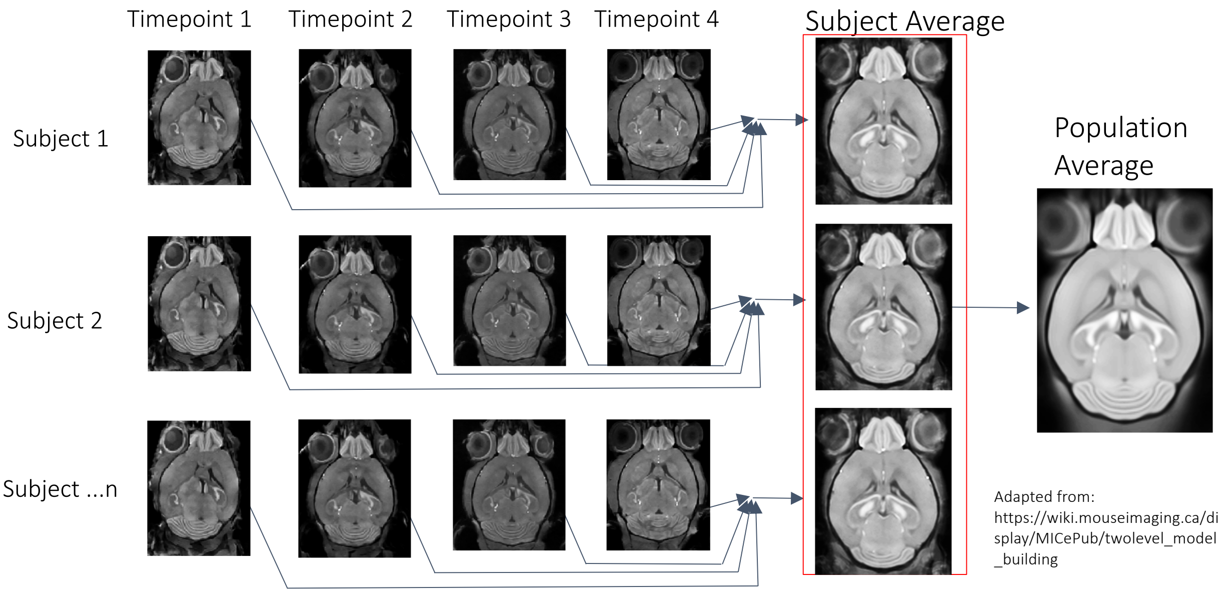


**Supplementary figure 1**. **Schematic of the registration pipeline**. Each subject (mouse) was scanned at 4 timepoints throughout development (timepoint 1, 2, 3, 4). The arrows indicate that the 4 scans for each subject were registered to create a subject average. All subject averages were then registered to create s population average for the entire study.

### 1.4. Behavioural testing

#### 1.4.1. Open field test

Mice were gently placed in the center of a 45 x 45 $cm^2$ rectangular light grey arena and allowed to explore for 15 minutes. Behaviour was video recorded. Areas were conceptually partitioned into zones in which distance traveled and time spent were measured: a center zone (40% of the total area), corners, and middle edges. We assessed distance traveled in the center zone relative to total distance traveled using the following model, which was applied to all tests and defined as behavioural model 1: group (SAL as reference) and sex (male as references) as fixed effects, and litter as a random effect to account for possible litter variability.

#### 1.4.2. Three chambered social preference and social novelty task

Mice were habituated (10 minutes) under red light to a three-chamber plastic box (26 (l) x 21.6 (w) x 21.6 (h) cm) with divider panels that have open doors, with a wire container (9.5 (h) 7.6 (d) cm) in each of the two extreme chambers. To measure social preference (10 minutes), time spent interacting with a stranger mouse was compared to that with a non social object using the following social preference index formula 1:

```
([distance traveled intruder 1 zone] / [distance traveled
object zone + distance traveled intruder 1 zone]) - 0.5.
(1)
```

Similarly, to measure social novelty (10 minutes), the nonsocial object was replaced with another stranger mouse, and a social novelty index was calculated as formula 2:

```
([distance traveled intruder 2 zone] / [distance
traveled intruder 1 zone + distance traveled intruder 2
zone]) - 0.5 (2)
```

We assessed group differences in social preference and novelty indices using behavioural model 1. Stranger mice were the same strain, sex, and similar age (within 2 weeks) of the test mice, and were habituated to the wire containers (20 minutes twice a day) for two days prior to the test.

#### 2.4.3. Marble burying task

Mice were placed in a standard home cage (28 (h) x 17 (w) x 12.7 (h) cm) filled with ~7 cm layer of woodchip bedding (new for each mouse) and 15 equidistant standard marbles for 30 minutes, as previously described (Deacon 2006). Marbles were classified as either buried 100% (nothing visible), buried at 75% (only a bit visible) or unburied (<75% buried). For the marble burying task, group differences in the number of marbles buried (75% and 100%) were tested with the behavioural model 1 described in the main text **2.4.2.** (group (SAL as reference) and sex (male as reference) as fixed effects, and litter as a random effect to account for possible variability due to litter ) using a Poisson distribution.

#### 1.4.2. Prepulse inhibition

Prepulse inhibition (PPI) to acoustic startle was measured using commercially available startle chambers (San Diego Instruments, San Diego, CA) consisting of a Plexiglass chamber (8 cm diameter, 16 cm long) mounted on a Plexiglass base with a sound-attenuating chamber, and a speaker located in the ceiling of the chamber (24 cm above the animal) to provide the background noise (70dB) and the acoustic stimuli. A piezoelectric accelerometer fixed to the animal enclosure frame was used to detect and transduce motion resulting from the animal's startle response. A microcomputer using a commercial software package by SR-LAB was used to control pulse parameters, and digitize (0-4095), rectify, and record the stabilimeter readings. Animals were placed in the

Plexiglass restrainers, and after 5 minutes of acclimatization. Mice underwent a total of 50 trials (5-30 s intertrial duration).

Startle magnitude to a 50 ms 120 dB stimulus, in absence of prepulse, was measured in the first 8 and final 7 trials. For the middle 35 trials, the startle tone was either presented alone, or preceded by a 30ms prepulse stimulus ranging from 3-15 dB above background noise (73-85 dB) and varying randomly between trials in 3 dB increments (5 trials per prepulse stimulus). A measure of maximum and average startle response was derived from the 100 1 ms readings taken starting from the beginning of the startle stimulus onset. Percent PPI from each prepulse intensity formula 3

```
(averaged over trials) was calculated using the formula:
[(startle response - prepulse response) / startle response]
x 100 (3)
```

Differences in maximum startle amplitude were investigated overall (behavioural model 1) and by prepulse intensity with a group * trial interaction added to behaviour model 1.

#### 1.4.3. Attentional set shifting task

***Food deprivation:*** four days following the PPI in adulthood, food deprivation was commenced for the attentional set shifting task (ASST). On the first day, the animal's body weight was measured at *ad libitum* conditions, and food was removed from all containers. Weight was maintained above 85% of initial body weight for each mouse by feeding 1 and 2 grams of food per mouse per day. One ceramic bowl was introduced per cage in which some pieces of Honey Nut Cheerios (General Mills, Canada) were placed as well as their regular chow in order to habituate mice to the bowls, and to reward food.

***Apparatus:*** testing chambers used for the ASST task consisted of opaque Plexiglass boxes (45 x 24x 18 cm) with white plastic walls (15 x 11.5 cm) to separate the two choosing compartments. Two ceramic bowls (6cm diameter) containing several digging mediums and odors were placed within each choice chamber. A separate plexiglass box was placed in front of the testing chamber to serve as a waiting area between trials. For each trial, mice were gently placed in the testing chamber with access to the two choice chambers.

***Habituation:*** after one day of food deprivation, mice were habituated to the testing chamber with two 20 minute sessions per day for four days. Mice were trained to find small pieces of Honey Nut Cheerio cereal (General Mills, Canada) by digging in two bowls containing bedding material. The amount of bedding medium covering the food reward was progressively increased on each day, until it was fully covered by the third day. Animals were required to dig efficiently (8 correct responses in a row), defined as vigorous displacement of medium with upper paws reaching the hidden reward in the training bowls, prior to starting ASST. Touching the rim or smelling the bowls at the surface of the medium was not considered digging.

***ASST Paradigm:*** the ASST comprised a successive series of discriminations in which mice had to choose between two bowls filled with different combinations of odors and digging mediums to find a hidden food reward. These were trained within 5 consecutive days and included simple discrimination (SD), compound discrimination (CD), intra-dimensional shift (ID), intradimensional shift reversal (IDR), and extradimensional shift (ED). Each stage was considered complete when mice reached a learning criterion of six consecutive correct responses. If a mouse stopped responding for three consecutive trials, the session was stopped, and training was resumed the following day (Colacicco et al. 2002). In the SD, one dimension (odor) was presented; mice had to discriminate between nutmeg (baited) and basil (unbaited). The next stage, CD, relied on the same previously trained odor discrimination (nutmeg/baited, basil/unbaited), however a new stimulus was introduced (medium: crystals and vase filler). For the ID, new combinations of odors (curry/baited, garlic/unbaited) and digging mediums (pompoms and shredded paper) were presented wherein mice were still required to discriminate according to the trained dimension: odor. For the IDR, the baited odor was switched from that of the previous day. Finally, for the ED, new pairs of stimuli were be introduced, but now the previously irrelevant stimulus dimension (medium) was relevant and predicted the baited bowl (wooden beads/baited and flower petals), whereas odor, the previously relevant stimulus now had to be ignored (paprika and ginger). The first four trials of the SD were exploratory in which mice were allowed to dig in the unbaited bowl and correct their choice (an error was still recorded); after this, mice were not allowed to correct their response.

**Supplementary table 1. Attentional Set Shifting Task (ASST) details**.

| | **Dimension** | | **Exemplar Combination** | | **Exemplar Compound Stimuli** | |
|---|---|---|---|---|---|---|
| **ASST Stage** | **Relevant** | **Irrelevant** | **Correct** | **Incorrect** | **Correct** | **Incorrect** |
| **SD** | Odor | Medium | O1, M1 | O2, M1 | Nutmeg/Bedding | Basil/Bedding |
| **CD** | Odor | Medium | O1, M2, M3 | O2, M2, M3 | Nutmeg/Vase Filler<br>Nutmeg/Crystals | Basil/Vase Filler<br>Basil/Crystals |
| **ID** | Odor | Medium | O3, M4, M5 | O4, M4, M5 | Curry/Shredded Paper<br>Curry/Pompoms | Garlic/Shredded Paper<br>Garlic/Pompoms |
| **IDR** | Odor | Medium | O4, M4, M5 | O3, M4, M5 | Garlic/Shredded Paper<br>Garlic/Pompoms | Curry/Shredded Paper<br>Curry/Pompoms |
| **ED** | Medium | Odor | M6, O5, O6 | M7, O5, O6 | Wooden beads/Ginger<br>Wooden beads/Paprika | Plastic flower petals/ginger<br>Plastic flower petals/parika |

**SD=** simple discrimination, **CD=** compound discrimination, **ID=** Intradimensional shift, **IDR=** Intradimensional shift reversal, **ED=** Extra-dimensional shift, **O=** odour, **M=** medium.

#### 1.5. Perfusions

One week following the last behavioural test (ASST), mice were anaesthetized with a pentobarbital overdose, and fixed by transcardiac perfusion with 4% paraformaldehyde (PFA) in phosphate buffered saline solution. Brains were collected and immersed in PFA overnight at 4 degrees Celsius for 24 hours, after which they were transferred to long-term storage solution of phosphate buffered solution with 0.02% Sodium Azide.

#### 1.6. Assessment of maternal cytokines levels

In a separate group of dams, poly I:C or saline was injected as described above (3 GD9-POL, 3 GD17-POL, 2 GD9-SAL, 2 GD17-SAL). Three hours following injection, dams were sacrificed by decapitation without euthanesia, and trunk blood was collected in a 1.5mL Eppendorf tube. The blood was allowed to coagulate at room temperature for 30 minutes, and then centrifuged for 10 minutes at 4 degree C, with 2000 revolutions per minute. Serum was collected and stored at -80 degrees C until ready for analysis. Serum samples were shipped to the University of Maryland Core Cytokine Facility (http://www.cytokines.com/) for multiplex ELISA to measure levels of IL-6, TNF-alpha, IL-1**β**, IL-10 to the immunostimulatory potential of our poly I:C. We chose to use a separate group of dams to ensure we could collect enough blood for analysis, and so as not to introduce an additional stressful experience for the dam, thereby potentially confounding the neurodevelopmental trajectory of offspring. Detection ranges were as follows IL-6 (1.95-8000 pg/ml), TNF-alpha (0.85-3500 pg/ml), IL-1**β** (3.75-15000 pg/ml), IL-10 (5-20000 pg/ml)

#### 1.7. Statistical analysis

##### 1.7.1.Model comparisons

Models were compared with increasing complexity using the minc Log Likelihood Ratio (LLR) function. This was computed for every voxel in the brain, therefore, for each compared model, we ensured that even if the model was significantly better based on LLR, it also covered a great proportion of the brain. We first tested what the best fit to model our age term was using non-linear splines for a linear fit (model 1), quadratic fit (model 2), or cubic fit (model 3).Next, inclusion of a random slope for age was also assessed (however the model did not converge, model 4). Finally, we investigated the effects of a 3 way interaction of age * group * sex (model 5).

**Model 1:** $Y_{subject,j} = \beta_0 + \beta_1 sex_{subject,j} + \beta_2 group_{subject,j} + \beta_3 age_{subject,j} + ns(\beta_4 age,1) : group_{subject,j} + v_{subject} + v_{litter} + \epsilon_{subject,j}$

**Model 2:** $Y_{ssubject,j} = \beta_0 + \beta_1 sex_{subject,j} + \beta_2 group_{subject,j} + \beta_3 age_{subject,j\ j} + ns(\beta_4 age,1) : group_{subject,j} + ns(\beta_4 age,2) : group_{subject,j} + v_{subject} + v_{litter} + \epsilon_{subject,j}$

**Model 3:** $Y_{ssubject,j} = \beta_0 + \beta_1 sex_{subject,j} + \beta_2 group_{subject,j} + \beta_3 age_{subject,j\ j} + ns(\beta_4 age,1) : group_{subject,j} + ns(\beta_4 age,2) : group_{subject,j} + ns(\beta_4 age,3) : group_{subject,j} + v_{subject} + v_{litter} + \epsilon_{subject,j}$

**Model 4:** $Y_{ssubject,j} = \beta_0 + \beta_1 sex_{subject,j} + \beta_2 group_{subject,j} + \beta_3 age_{subject,j\ j} + ns(\beta_4 age,1) : group_{subject,j} + ns(\beta_4 age,2) : group_{subject,j} + ns(\beta_4 age,3) : group_{subject,j} + age|v_{subject} + v_{litter} + \epsilon_{subject,j}$

**Model 5:** $Y_{ssubject,j} = \beta_0 + \beta_1 sex_{subject,j} + \beta_2 group_{subject,j} + \beta_3 age_{subject,j\ j} + ns(\beta_4 age,1) : sex:group_{subject,j} + ns(\beta_4 age,2) : sex:group_{subject,j} + ns(\beta_4 age,3) : sex:group_{subject,j} + v_{subject} + v_{litter} + \epsilon_{subject,j}$

**Y**= outcome measures (i.e. blurred absolute Jacobian determinants); $\boldsymbol{\beta}_i$: fixed effect coefficient; $\boldsymbol{\beta}_0$ = equation intercept; **v** = random intercept; ϵ: random error; $_j$ = repeated measure (timepoint)

#### 1.7.2. Comparison of SAL E and SAL L

We investigated whether there were differences in neuroanatomical development in our two control groups (SAL E and SAL L). Using a brain mask, we ran a whole-brain voxel-wise linear mixed effects model on the subject-level (first level) absolute Jacobian determinant files for each subject over all 4 timepoints. We assessed an age, modeled as a cubic spline, by injection timing interaction, with sex as a covariate (fixed effects), and used mouse id and litter as random intercepts. A False Discovery Rate (FDR) correction was applied to correct for multiple testing.

#### 1.7.3. Partial Least Squares Analysis

***Inputs:*** PLS cannot handle missing data. Thus, in order to maximize our ability to detect brain-behaviour patterns, we performed data imputation on the behavioural data so that any mouse with a scan passing QC could have a full set of behavioural metrics to use (22 mice were removed). We did this using the singular value decomposition function SVD.miss in R (Fuentes et al. 2006). Briefly, this completes a data matrix using iterative singular value decomposition, replacing the missing values by linear regression of the columns, and making use of the column averages. The input brain matrix included whole-brain voxel-wise DBM measures (87 by 508169 voxel matrix) from the PND38 scan. The imputed input behavioural data included behavioural metrics for all tests performed immediately following that PND38 scan (78 x 20 matrix including: OFT: velocity in center, total distance, distance moved in center; SOPT: frequency of entries in object circle, and intruder 1 circle, ratio of time spent in intruder 1 vs. object circle, ratio of distance moved in intruder 1 and object boxes; SONT: same as SOPT; PPI: PP6 max, PP9 max, PP12 max, PP15max; marbles buried 100%, marbles buried 75%). As described in the main text, sex and litter size were also included in the behaviour matrix as 'demographic' measures. The imaging and behavioural data were organized into two matrices X (imaging), and Y (behaviour), with subjects as rows and variables in columns.

***Permutation testing:*** was used to assess the statistical significance of each LV wherein the rows (subjects) of the brain data matrix were randomly shuffled to 1) nullify dependencies between brain and behaviour (n=1000 repetitions) and 2) generate a null distribution of possible brain-behaviour correlations. SVD was applied to these "null" correlations, generating a distribution of singular values under the null hypothesis. The probability that a permuted singular value exceeds the original, non-permuted singular value allows us to generate the p-value (Zeighami et al. 2019; Patel et al. 2020). A threshold of p<0.05 was used (95% or greater chance that the singular value of the non permuted data exceeds that of a permuted singular value).

***Bootstrap resampling:*** was applied to assess the contribution of individual brain and behaviour variables to each LV. Subjects (rows for both X and Y matrices) were randomly sampled and replaced (n=1000) to generate a set of resampled correlation matrices to which SVD was applied to generate a sampling distribution for each weight of the singular vectors. The ratio of each singular vector weight and its bootstrap-estimated standard error were used to calculate a "bootstrap ratio" for each voxel. Voxels that make large contributions to certain patterns can therefore be identified by large bootstrap ratios.

### 1.8. Transcriptional analysis

#### 1.8.1. Pre-processing

Sequencing was performed with Illumina NovaSeq 6000 for 72 samples, 3 regions in adolescent POL E or SAL E mice (PND38; 6males/6females per group). Pre-alignment quality control was performed using FastQC (version, reference).

Reads were filtered for a minimum Phred score of 30 and a minimum read length of 20 as well as trimmed off the first base pair from the 5' end. Alignment was performed using the STAR aligner (Version 2.7.3a_2020-01-23) (Dobin et al. 2013) on the mouse genome build (mm10)/(GRCm38.p6)(GCA_000001635.8) downloaded from Ensembl

Non-specific filtering removed genes with zero counts and lowly expressed genes that did not meet the requirement of a minimum of one count per million (cpm) in at least six samples. Only genes annotated as protein coding according to ensembl's biomart Mus.musculus package (https://bioconductor.org/packages/release/data/annotation/html/Mus.musculus.html) were retained (15048 genes). Genes were subjected to a trimmed mean of M-values normalization method (TMM) (Robinson and Oshlack 2010). Normalized data were inspected for outlier samples using unsupervised hierarchical clustering of subjects by multidimensional scaling (MDS) and principal component analysis (PCA) to identify potential outliers greater than two standard deviations from these averages. Outlier detection was also performed using high dimensional extension of Cook's influence measure (Cook 1986), which identified no outliers, however one sample (ACC, F, TX) was removed as it was close to the outlier cutoff and displayed expression signals in Y chromosome genes (indicative of possible contamination).

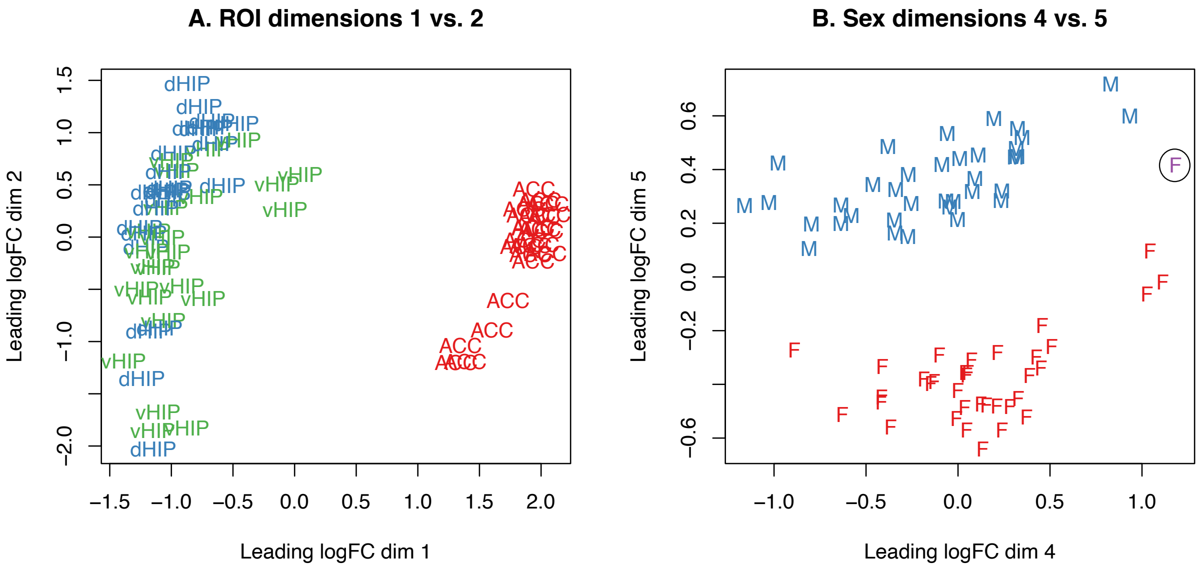


**Supplementary figure 2.** Multidimensional scaling (MDS) plots of log-counts per million (CPM) Values. **A.** Over dimensions 1 and 2 with samples coloured and labeled by region of interest. **B** Over dimensions 4 and 5 with samples coloured and labeled by sex. Distances on the plot correspond to the leading fold-change, which is the average (root-mean-square) log2-fold-change for the 500 genes most divergent between each pair of samples by default. Magenta circle denotes the sample that was excluded from analysis (**ACC**=anterior cingulate cortex; **dHIP**=dorsal hippocampus; **vHIP**=ventral hippocampus; **M**=male; **F**=female; **ROI**=region of interest).

#### 1.8.2. Differential Expression Analysis

DE analysis was performed according to the limma-voom pipeline (Ritchie et al. 2015), using a generalized linear model (GLM), log-normal distribution and a nominal significance threshold multiple testing adjusted p-values of $p<0.05$. Correction for multiple comparisons was performed using FDR (Benjamini and Hochberg 1995). The voomWithQualityWeights() method (Liu et al., 2015) was used to assign additional weight to more variable samples, allowing for more statistical power and accounting for heteroscedastic nature of the data. Sex and ROIs were used as confounding variables for the GLM model as well as blocking on the individual level using limma's duplicateCorrelation() method (Smyth et al., 2005), to account for the 3 ROIs being taken from the same specimen. T-statistics, moderated F-statistic, and log-odds of differential expression were estimated by limma's ebayes() method using the robust parameter to account for individual gene outliers (Phipson et al., 2016).

#### 1.8.3 Rank Rank Hypergeometric Overlap Test

We applied a rank rank hypergeometric overlap test (RRHO; recently updated version of the software (Cahill et al., 2018)), which builds on the previous version by accurately detecting overlap of the change in genes in both the same and opposite directions in the same dataset, to measure the concordance of differential gene expression patterns between POL E and SAL E data in a multitude of settings. RRHO is a threshold-free approach to identify concordant and discordant overlaps between expression profiles and to measure the degree and significance of said overlap (Plaisier et al., 2010). Full differential expression lists were ranked by the -log10(p-value) multiplied by the sign of the fold change from the DE analysis. We assessed concordance between the sexes (male (M) vs. female (F)) for all brain regions combined as well as within each individual brain region. RRHO maps compute the normal approximation of difference in log odds ratio and standard error of overlap between conditions POL vs. SAL (control) for each pixel. This Z score is then converted to a P-value and corrected for multiple comparisons across pixels (Benjamini and Yekutieli, 2001).

#### 1.8.4 Pathway enrichment analysis

In order to gain further insight from the DE and RNA-Seq results, we performed pathway enrichment analysis. Firstly, we used the g:Profiler (Raudvere et al., 2019) R client to identify pathways whose genes are significantly enriched or overrepresented in a gene list of interest compared to a background gene list. Next, we performed Gene Set Enrichment Analysis (GSEA) (Subramanian et al., 2005) on full ranked gene lists of DE results. Resulting pathways were selected using an FDR adjusted p value threshold (Q value) $< 0.05$ and ranked by normalized enrichment score (NES). Finally, to organize and visualize pathway analyses results, we used the EnrichmentMap (Merico et al., 2010) and Autoannotate (Kucera et al., 2016) packages made available within the Cytoscape platform (Shannon et al., 2003).

#### 1.8.5 Gene Overlap Analysis

To compared our resulting differentially expressed gene (DEG) lists to published data on human diseases, notably schizophrenia (SCZ) DEGs (Lanz et al. 2019) and SCZ and autism spectrum disorder (ASD) pancortical DEGs (Gandal et al. 2018), we used the GeneOverlap package (Li Shen and Mount Sinai (2019), R package version 1.20.0.) to probe enrichment of disease gene lists in our DEG lists by Fisher's exact test..

2. **Results**

#### 2.1. Poly I:C injection does increase pro-inflammatory cytokines

We observed an increase in levels of pro-inflammatory cytokines IL-6 and IL-1**β**, but not TNF-α, in a separate cohort of pregnant dams 3-hours post poly I:C injection on GD9 relative to saline control on GD9. Exposure to poly I:C on GD17 increased levels of pro-inflammatory cytokines IL-6 and TNF-α, but not IL-1**β** relative to GD17 saline controls. There were no differences in levels of anti-inflammatory cytokine IL-10 for any of the 4 groups.

**Supplementary Table 2. Maternal serum cytokine levels for our 4 treatment groups, mean [range]**

| | **IL-1β (pg/ml)** | **IL-6 (pg/ml)** | **IL-10 (pg/ml)** | **TNF-α (pg/ml)** |
|---|---|---|---|---|
| **SAL E (n=2)** | 5.00 [3.65-6.34] | 32.95 [30.58-35.32] | 33.83 [11.43-56.22] | 23.39 [20.72-26.05] |
| **SAL L (n=2)** | 6.610 [6.34-6.88] | 23.27 [22.22-24.31] | 21.52 [8.30-34.73] | 6.84 [5.99-7.69] |
| **POL E (n=3)** | 11.56 [9.80-12.95] | 3578.84 [3624.87-3740.81] | 44.05 [36.93-49.70] | 8.20 [7.91-8.35] |
| **POL L (n=3)** | 5.45 [5.00-6.34] | 1298.69 [54.36-3173.25] | 16.73 [13.67-19.58] | 20.35 [14.21-31.18] |

#### 2.2. Model comparisons

Using the minc log likelihood function (LLR) in R, we found that model 3 (cubic nonlinear spline fit for age) was better than models 1 (linear) and 2 (quadratic) (LLR=13.702141, q=0.01). The addition of sex as an interaction term did not result in a better fitting model (model 5), and the addition of a random slope for age was too complex of a model to fit, as it did not converge (model 4). Thus, we chose model 3 for our analyses.

A.

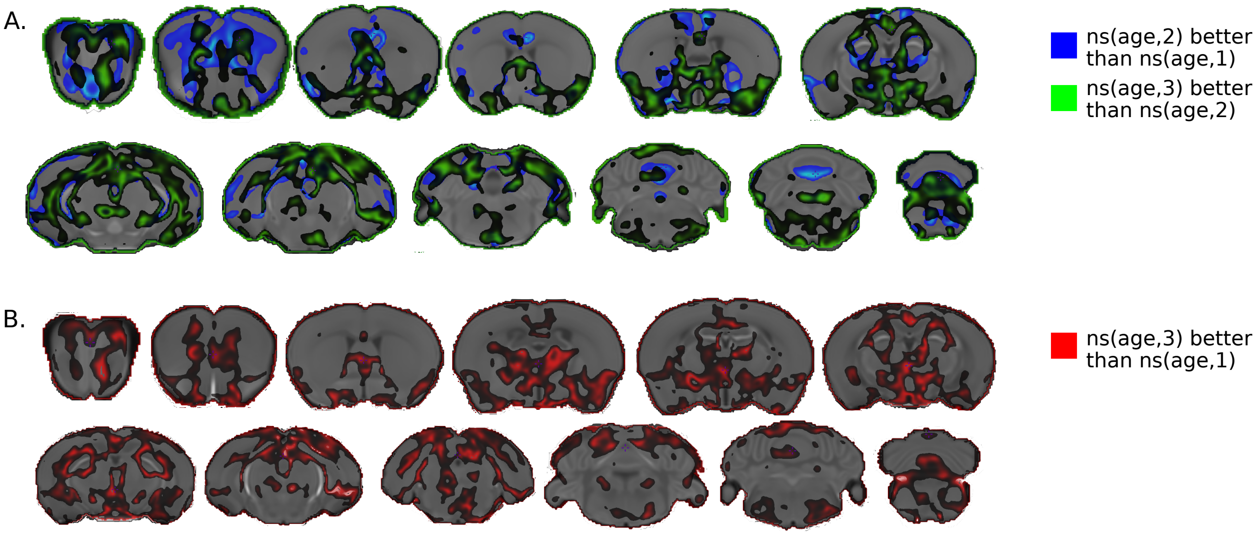


**Supplementary figure 3.** Comparison of 3 linear mixed-effects models with different age fits. **A.** Regions in blue were better fit by modeling age as a quadratic natural spline (ns(age,2)) relative to a linear age fit (ns(age,1)). Whereas regions in green were better modeled by a cubic natural spline fit for age (ns(age,3)) relative to the quadratic fit (ns(age,2)) according to the log-likelihood ratio test ($q<0.05$). **B.** Regions in red were better modeled by a cubic natural spline fit (ns(age,3)) relative to a linear fit (ns(age,1)) ($q<0.05$), supporting our choice of a cubic fit. Maps are overlaid on the population average from the study.

**2.3. Longitudinal neuroanatomical changes due to early and late MIA-exposure for all age fits: linear, quadratic, cubic.**

We present a more detailed results figure for regions affected by early MIA-exposure for the quadratic age term, and late MIA-exposure for the cubic age term, both highlighted in the main text.

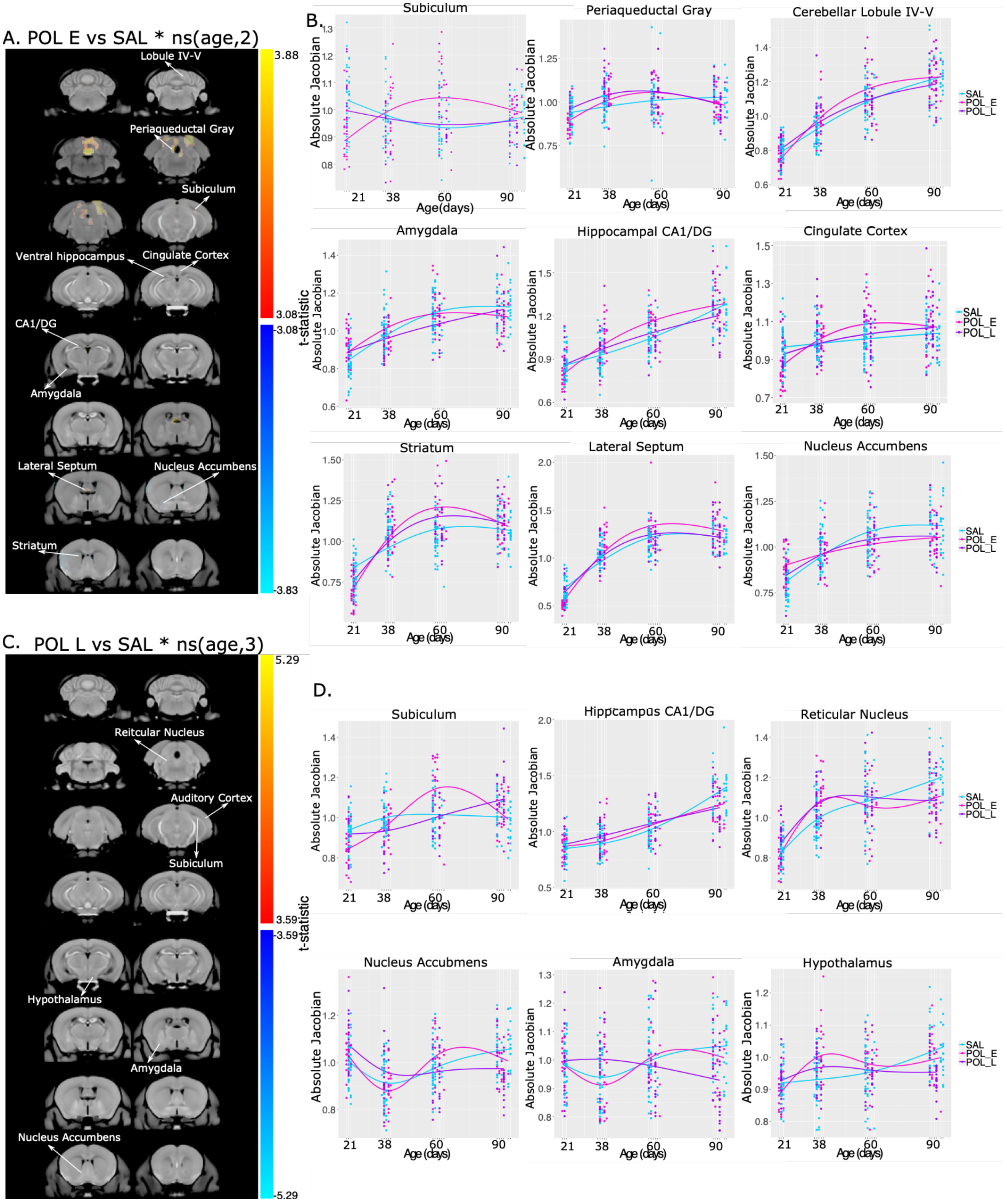


**Supplementary figure 4**. Differences in developmental trajectories for early poly I:C group (POL E) vs saline controls (SAL) & the late poly I:C group (POL L) vs SAL (thresholded at 5% False discovery rate (FDR)) (**A**) t-statistic map of group (POL E vs SAL) by age (quadratic natural spline)

thresholded between 5% FDR (bottom, t=3.08) and 1% FDR (top, t=3.83) overlaid on the population (second level) average (**B**) Plot of peak voxels (voxel within a region of volume change showing largest effect) selected from regions of interest highlighted (**A**), wherein age is plotted on the x-axis, and the absolute Jacobian determinants plotted on the y-axis. Here a value of 1 means the voxel is no different than the average, anything above 1 is relatively larger, and below 1 is relatively smaller. Ranges are not normalized to enhance comparison at each specific location in space. Trajectories are modeled as quadratic natural splines to reflect statistical modeling. (**C**) t-statistic map of group (POL L vs SAL) by age (cubic natural spline) thresholded between 5% FDR (bottom, t=3.59) and 1% FDR (top, t=5.29). (**D**) Plots of peak voxels as described in (**B**) with curves modeled as cubic natural splines to reflect statistics.

The interaction between cubic age and group was significant for POL E offspring relative to SAL (t= 4.323, <5%FDR) in small subregions of the right and left subiculum, auditory, motor, and posterior cingulate cortex. This was reflective of an overgrowth in the early adult period in the POL E offspring. Finally, the linear age by group interaction was also significant in many regions for POL E vs. SAL offspring (t=3.035, <1%FDR), indicative of a steeper increase in volume over time for POL E offspring relative to the SAL offspring. These regions are mainly cortical, including the motor and somatosensory cortices, visual, and auditory, as well as the bilateral striatum, thalamus, hypothalamus with some subregions of the dorsal and ventral hippocampus as well.

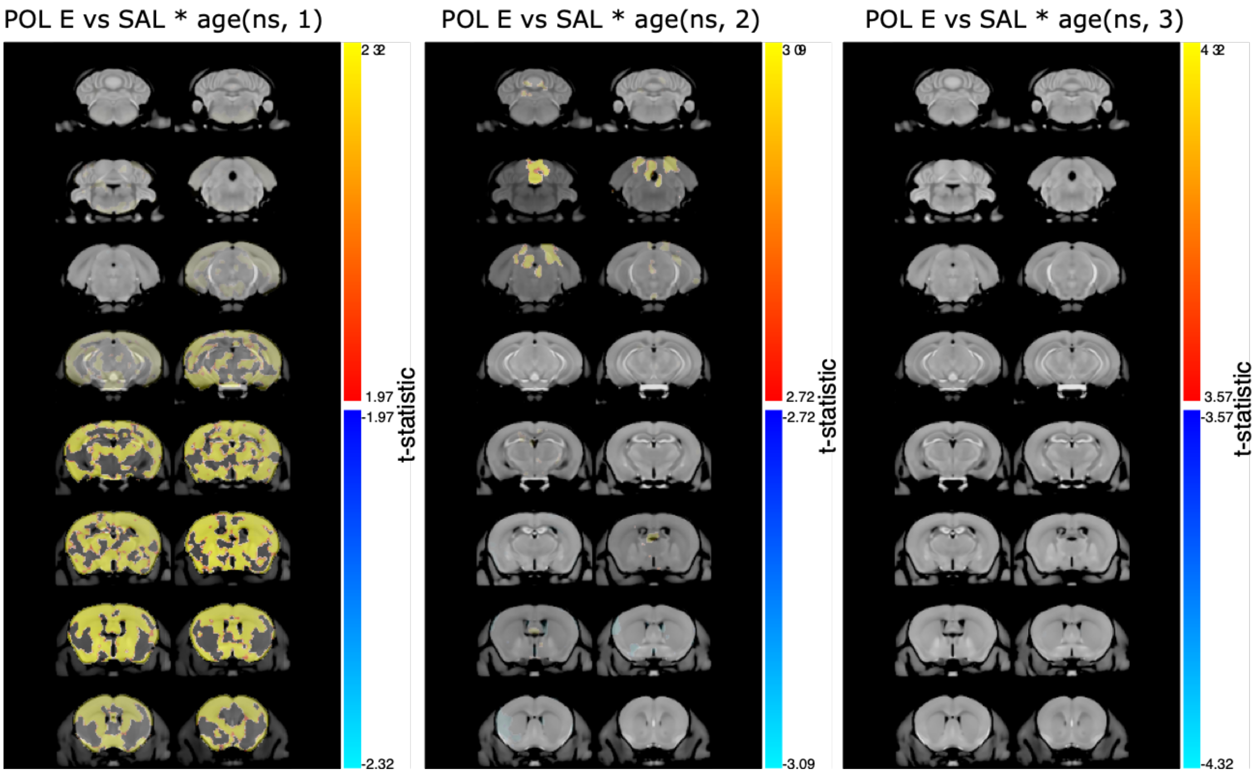


**Supplementary figure 5.** T-statistic maps for POL E vs SAL linear age (left), quadratic age (middle), and cubic (right) age overlaid on population average (t-statistics thresholded between 10% and 5% FDR).

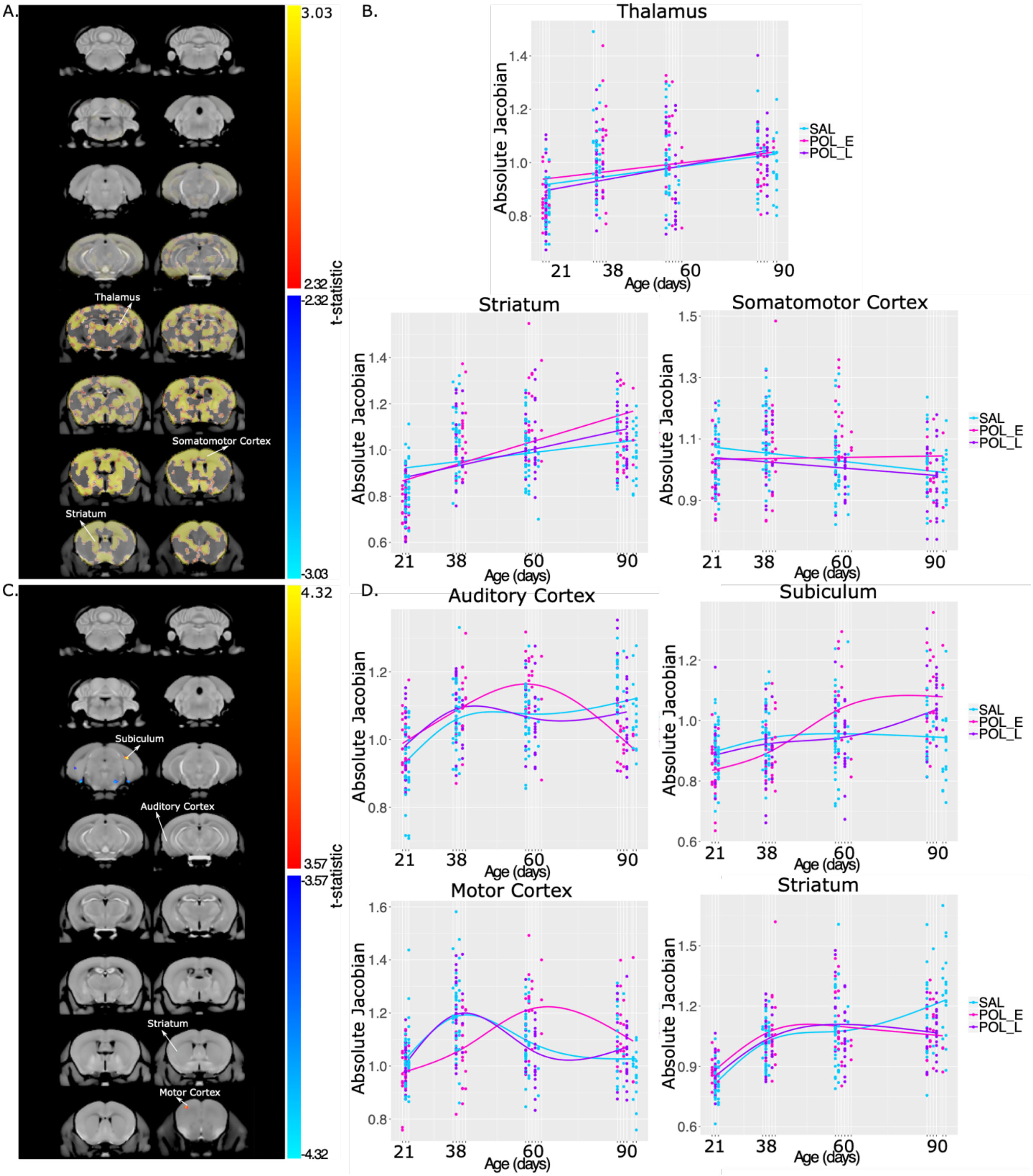


**Supplementary figure 6.** POL E vs SAL linear age and cubic age plots. (**A**) t-statistic map of group (POL E vs SAL) by age (linear natural spline) (t=3.035, <1%FDR). (**B**) Plot of peak voxels selected from regions of interest highlighted (**A**), wherein age is plotted on the x-axis, and the absolute Jacobian determinants plotted on the y-axis. Here a value of 1 means the voxel is no different than the average, anything above one is relatively larger, and blow 1 is relatively smaller. Ranges are not normalized to enhance comparison at each specific location in space. Lines are modeled as quadratic natural splines to reflect statistics. (**C**) t-statistic map of group (POL E vs

SAL) by age (cubic natural spline) (t= 4.323, <5%FDR). (**D**) Plots of peak voxels as described in (**B**) with curves modeled as cubic natural splines to reflect statistics.

#### 2.4. Longitudinal comparison of POL L vs SAL for quadratic and linear age.

The interaction between quadratic age and POL L relative to SAL was significant (t=4.523, <1%FDR) in the bilateral ventral tegmental area, the striatum, and tenia tecta. In these regions, the POL L offspring displayed a flatter curve relative to SAL (and POL E). Finally, the interaction between POL L and linear age was statistically significant (t=4.519, <5%FDR) only in a few negligible voxels.

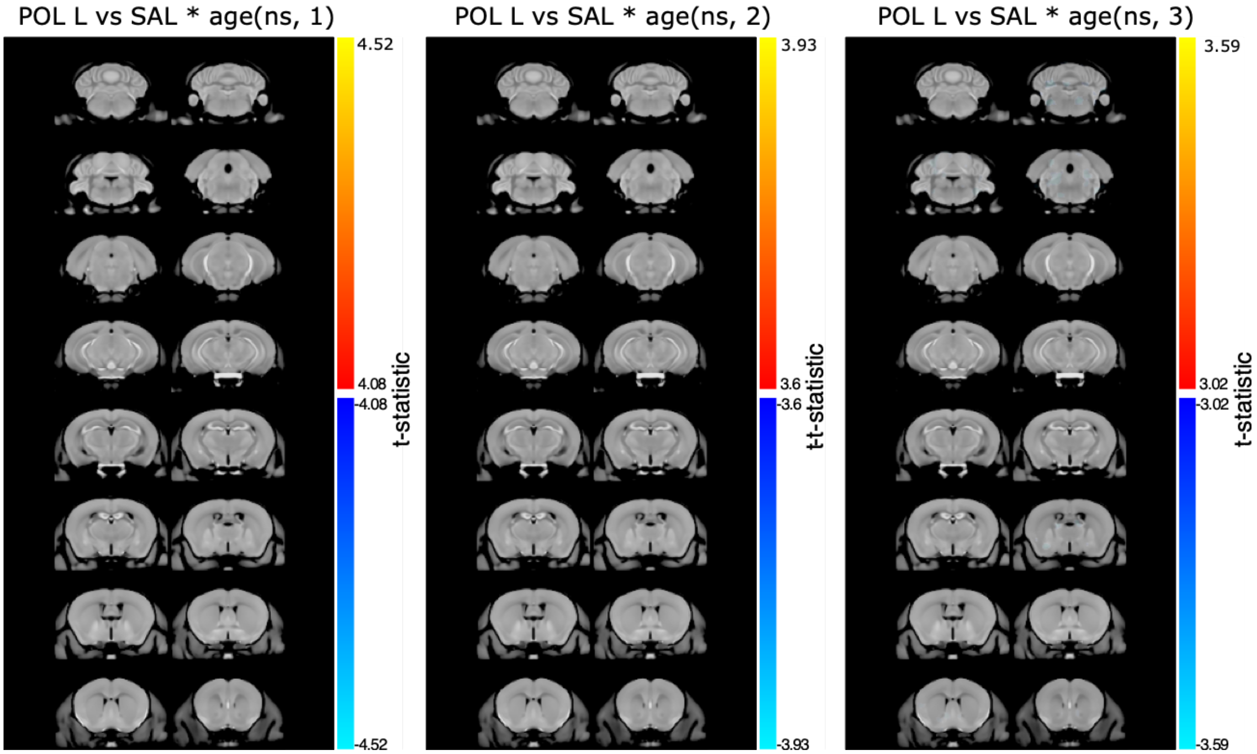


**Supplementary figure 7.** T-statistic maps for POL L vs SAL linear age (left), quadratic age (middle), and cubic (right) age overlaid on population average (t-statistics thresholded between 5% and 10% FDR).

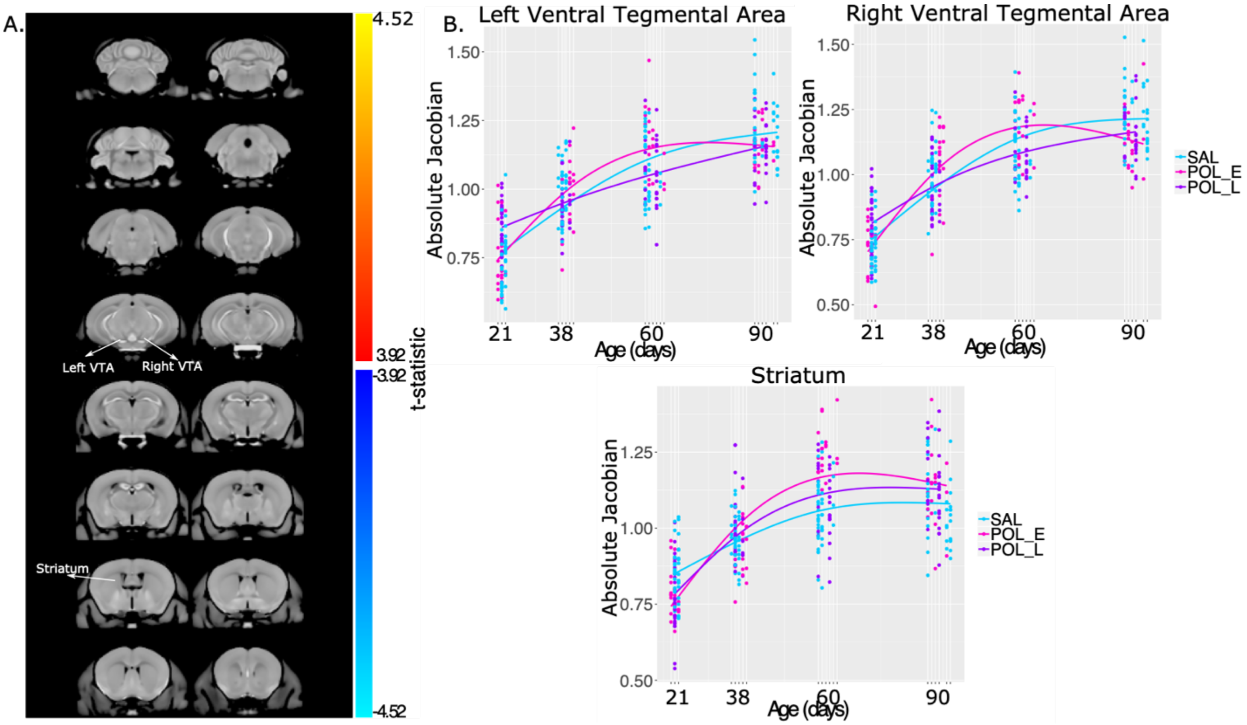


**Supplementary figure 8.** POL L vs SAL quadratic age plots. (**A**) t-statistic map of group (POL L vs SAL) by age (quadratic natural spline) (t=4.523, <1%FDR). **B**) Plot of peak voxels selected from regions of interest highlighted (**A**), wherein age is plotted on the x-axis, and the absolute Jacobian determinants plotted on the y-axis. Lines are modeled as quadratic natural splines to reflect statistics.

### 2.5. Longitudinal comparison of POL E vs. POL L

We re-ran our statistical analysis with the POL L group as the reference group rather than SAL in order to better compare differences between the two POL exposed groups. We observed a significant group (POL E vs. POL L) by cubic age interaction (t=4.268, <10%FDR) in a few voxels in the amygdala, nucleus accumbens, thalamus, hippocampus. There was also a significant group (POL E vs POL L) by quadratic age (t=3.977, <1%FDR) in very similar regions to what was observed for the POL E vs SAL comparison, including the the striatum, lateral septum, CA1 and dentate gyrus of the hippocampus, subiculum, thalamus, periaqueductal gray, and cerebellum; additionally differences in the VTA/substantia nigra, bed nucleus of stria terminalis were also observed. These regions displayed a similar trajectory as described in the POL E vs SAL comparison whereby the POL E group had a smaller volume at PND21, followed by an overshoot in the P38-60 period, and a normalization at PND 90. Finally, there was also a significant effect for the interaction with linear age in the majority of the cortex, as well as the striatum, and many thalamic regions, indicative of a steeper growth in the POL E relative to the POL L group (t=3.001, <1%FDR).

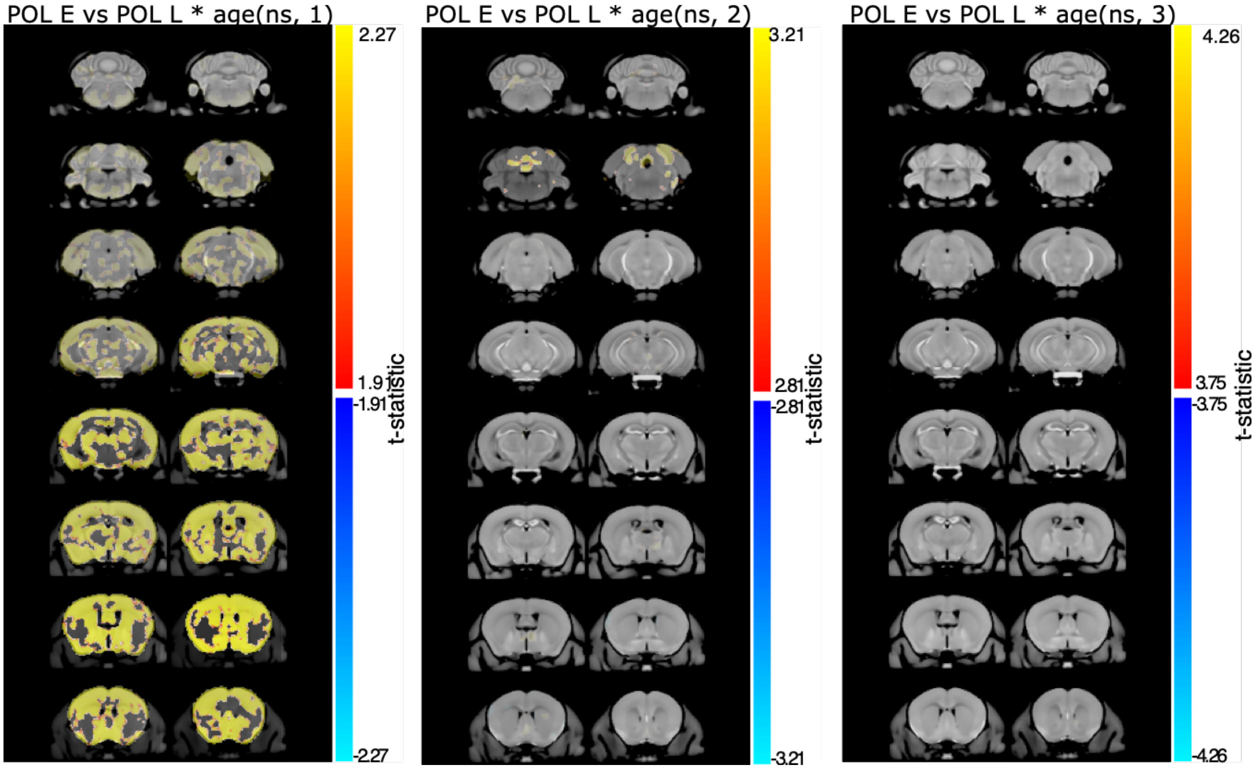


**Supplementary figure 9.** T-statistic maps for POL E vs POL L linear age (left), quadratic age (middle), and cubic (right) age overlaid on population average (t-statistics thresholded between 5% and 10% FDR).

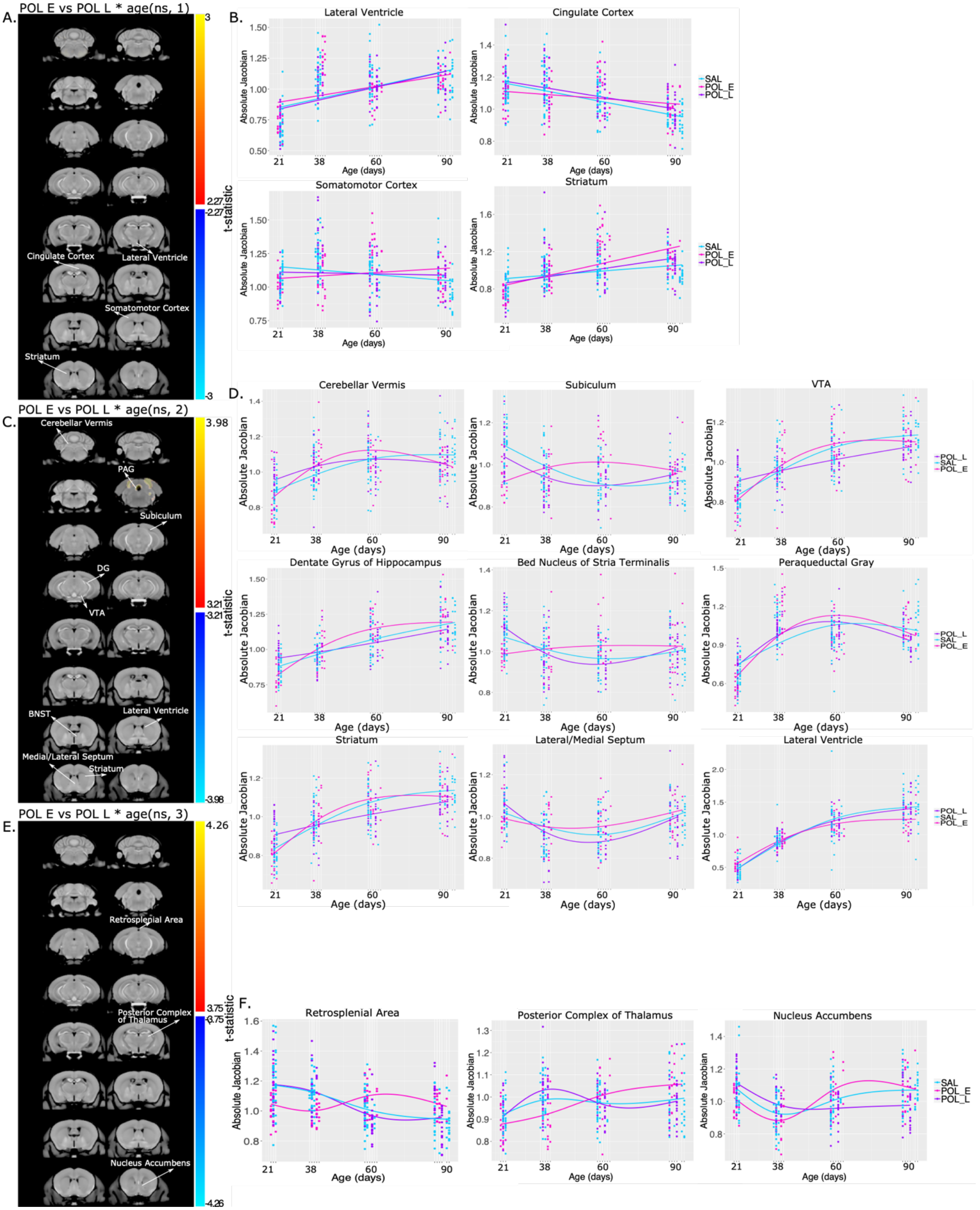
A.
POL E vs POL L * age(ns, 1)
Cingulate Cortex
Lateral Ventricle
Somatomotor Cortex
Striatum
t-statistic
B.
Lateral Ventricle
Cingulate Cortex
Somatomotor Cortex
Striatum
Absolute Jacobian
Age (days)
SAL
POL_E
POL_L
C.
POL E vs POL L * age(ns, 2)
Cerebellar Vermis
PAG
Subiculum
DG
VTA
BNST
Lateral Ventricle
Medial/Lateral Septum
Striatum
D.
Cerebellar Vermis
Subiculum
VTA
Dentate Gyrus of Hippocampus
Bed Nucleus of Stria Terminalis
Peraqueductal Gray
Striatum
Lateral/Medial Septum
Lateral Ventricle
E.
POL E vs POL L * age(ns, 3)
Retrosplenial Area
Posterior Complex of Thalamus
Nucleus Accumbens
F.
Retrosplenial Area
Posterior Complex of Thalamus
Nucleus Accumbens

**Supplementary figure 10.** POL E vs POL L linear age, quadratic age, and cubic age plots. (A) t-statistic map of group (POL E vs POL L) by age (linear natural spline) (t=3.001, <1%FDR). B) Plot of peak voxels selected from regions of interest highlighted (A), wherein age is plotted on the x-axis, and the absolute Jacobian determinants plotted on the y-axis. Lines are modeled as linear natural splines to reflect statistics. (C) t-statistic map of group (POL E vs POL L) by age (quadratic natural spline) thresholded (t=3.977, <1%FDR). (D) Plots of peak voxels as described in (B) with curves modeled as quadratic natural splines to reflect statistics. (E) t-statistic map of group (POL E vs POL L) by age (cubic natural spline) (t=4.268, <10%FDR). (F). Plots of peak voxels as described in (B) with curves modeled as cubic natural splines to reflect statistics.

### 2.6. Longitudinal sex differences

Our longitudinal analysis revealed a no significant sex by group by age interactions with SAL as the reference group. However, with POL L as the reference group, there was a significant POL E by sex by age interaction for the quadratic age fit (t=3.980, <5%FDR). In regions such as the dorsal striatum, lateral septum, orbital cortex, dentate gyrus, hippocampal CA3, cingulate cortex, and parieto-visual cortex, male offspring seemed to have a more pronounced volume peak in the adolescent/early adult period relative to POL L offspring, whereas the difference between female offspring was much more subtle. Similarly, the interaction for linear age was also significant, albeit at a more lenient threshold (t=4.907, <10%FDR), in the left striatum, wherein the male offspring also had a more pronounced volume increase over time than females.

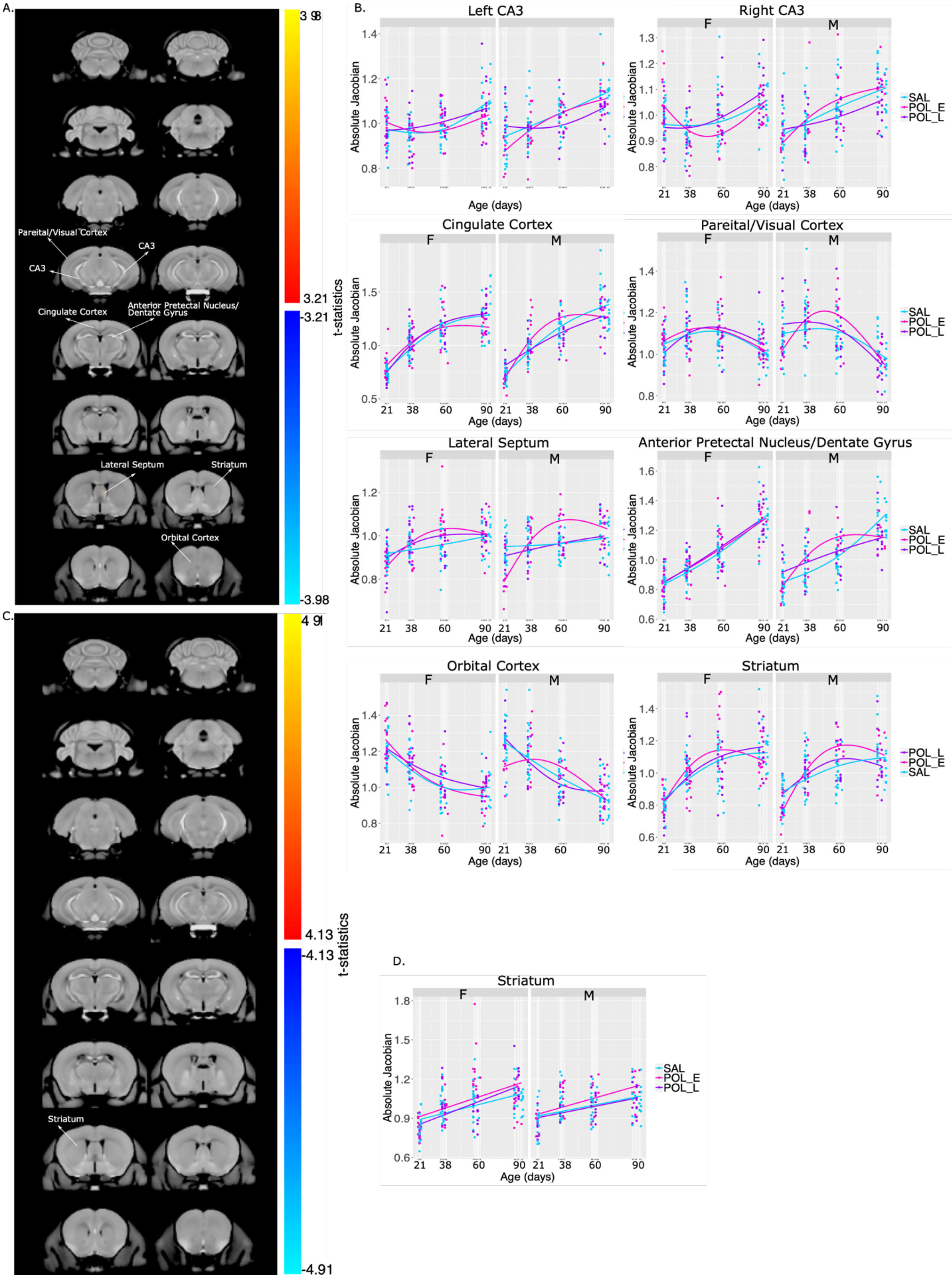
A.
Pareital/Visual Cortex
CA3
CA3
Cingulate Cortex
Anterior Pretectal Nucleus/
Dentate Gyrus
Lateral Septum
Striatum
Orbital Cortex
3.21
-3.21
-3.98
t-statistics
B.
Left CA3
Right CA3
Cingulate Cortex
Pareital/Visual Cortex
Lateral Septum
Anterior Pretectal Nucleus/Dentate Gyrus
Orbital Cortex
Striatum
F
M
Absolute Jacobian
Age (days)
SAL
POL_E
POL_L
C.
Striatum
4.13
-4.13
-4.91
t-statistics
D.
Striatum
F
M
Absolute Jacobian
Age (days)
SAL
POL_E
POL_L

**Supplementary figure 11.** Sex differences in brain development between POL E and POL L offspring. (A) T-statistic map of group (POL E vs. POL L) by age (quadratic natural spline) by sex interaction thresholded between 5% FDR (t=3.980). (B) Plot of peak voxels selected from regions of interest highlighted (A), wherein age is plotted on the x-axis, and the absolute Jacobian determinants plotted on the y-axis.(C) t-statistic map of group (POL E vs POL L) by age (linear natural spline) thresholded at 10% FDR (bottom, t=4.91) (D) Plots of peak voxels as described in (B) with curves modeled as linear natural splines to reflect statistics.

### 2.7. Behavioural findings

#### 2.7.1. Main effects

##### 2.7.1.1. Social Preference and Social Novelty

Prenatal MIA exposure did not affect significantly affect preference for a novel mouse over inanimate object for either adolescent POL E (t=-1.109, p=0.289) or POL L (t=-0.209, p=0.839) offspring or adult POL E (t=1.818, p=0.093) or POL L offspring (t=0.380,p=0.7123). Similarly, there were no significant group differences for the social novelty portion of the task for adolescent POL E (t=-0.330, p=0.742) or POL L (t=0.447, p=0.656) offspring. In adulthood, there was a significant impairment in social novelty behaviour for POL E offspring (t=-2.369, p=0.0311) wherein they spent less time exploring the novel intruder relative to the familiar one. Adult POL L offspring did not differ significantly from SAL offspring (t=-1.257, p=0.233).

##### 2.7.1.2. PPI

Adolescent POL E offspring were found to have significant impairment in sensorimotor gating, with significantly decreased percent prepulse inhibition relative to SAL (t=-4.335, p=0.0000005). No difference was observed for POL L offspring relative to SAL (t=-0.373, p=0.709). In adulthood, no differences were observed for POL E (t=-0.283, p=0.7837) or POL L (t=-0.549, p=0.5983) offspring.

We also investigated whether or not there was an interaction between increasing prepulse level and group. We did not observe any significant differences in trajectory in adolescence for POL E (t=-0.995, p=0.3209) or POL L (t=-0.887, p=0.376361) offspring in adolescence. We did observe a significant interaction for POL L offspring in adulthood (t=-2.211, p=0.0281) but not for POL E offspring (t=-1.401, p=0.163). There was a significant interaction between prepulse level and POL L group (t=-2.211, p=0.0281) indicative of POL L deficits emerging only at louder prepulse tones (12 and 15 dB). Finally, on the ASST, adult POL L offspring displayed impaired learning and flexibility as they required significantly more trials than SAL to reach learning criterion on the intradimensional shift portion of the task (t=2.486, p=0.0129).

**Table 3. Summary of all behavioural results for POL E and POL L offspring relative to SAL controls.** P-values are uncorrected, but only bolded if they survive Bonferroni correction (q-value=p<0.0045).

| | **Adolescence** | | **Adulthood** | |
|---|---|---|---|---|
| | **POL E** | **POL L** | **POL E** | **POL L** |
| **OFT** | t=-2.294, p=0.039, q=0.429 | t=-1.716, p=0.108, q=1.188 | t=1.116, p=0.281, q=3.091 | t=-0.069, p=0.946, q=10.406 |
| **Marble Burying** | **t=2.937, p=0.003, q=0.033** | t=0.901, p=0.368, q=4.048 | t=1.055, p=0.291, q=3.201 | t= 0.117, p=0.907, q=9.977 |
| **SOPT** | t=-1.109, p=0.289, q=3.179 | t=-0.209, p=0.839, q=9.229 | t=1.818, p=0.093, q=1.023 | t=0.380, p=0.712, q=7.832 |
| **SONT** | t=-0.330, p=0.742, q=8.162 | t=0.447, p=0.656, q=7.216 | t=-2.369, p=0.031, q=0.341 | t=-1.257, p=0.233, q=2.563 |
| **PPI** | Overall: **t=-4.202, p=0.0000004, q=0.000004** By PP tone: t=-0.995, p=0.321, q=3.531 | Overall: t=-0.373, p=0.709, q=7.799 By PP tone: t=-0.887, p=0.376, q=4.136 | Overall: t=-0.283, p=0.784, q=8.624 By PP tone: t=-1.401, p=0.163, q=1.793 | Overall: t=-0.549, p=0.598, q=6.578 By PP tone: t=-2.211, p=0.0281, q=0.310 |
| **ASST** | NA | NA | CD: t=-0.823, p=0.411, q=5.521 ID:t=0.522, p=0.602, q=6.622 IDR:t=0.918, p=0.359, q=3.949 ED:t=-1.879, p=0.060, q=0.660 | CD:t=1.853, p=0.064, q=0.704 ID:t=2.486, p=0.0129, q=0.142 IDR: t=0.158, p=0.874, q=9.614 ED: t=0.535, p=0.593, q=6.523 |

**POL E**= GD9-exposed poly I:C group; **POL L**=GD17-exposed poly I:C group; **OFT**= open field test; **SOPT=** social preference task; **SONT**= social novelty taks; **PPI**= prepulse inhibition; **ASST**= attentional set shifting; **SD=** simple discrimination, **CD=** compound discrimination, **ID=** Intradimensional shift, **IDR=** Intradimensional shift reversal, **ED=** Extra-dimensional shift.

##### 2.7.1. ASST

Adult POL L offspring required more trials that SAL to reach learning criterion on the IDR portion (t=2.486, p=0.0129) suggesting some impairment in learning; they also required slightly more trials on the CD portion of the task ((t=-1.853 0.064). POL E offspring required fewer trials than SAL in the ED portion of the task (t=-1.879, p=0.0602), perhaps indicative of greater flexibility or poorer learning of the previous day's task.

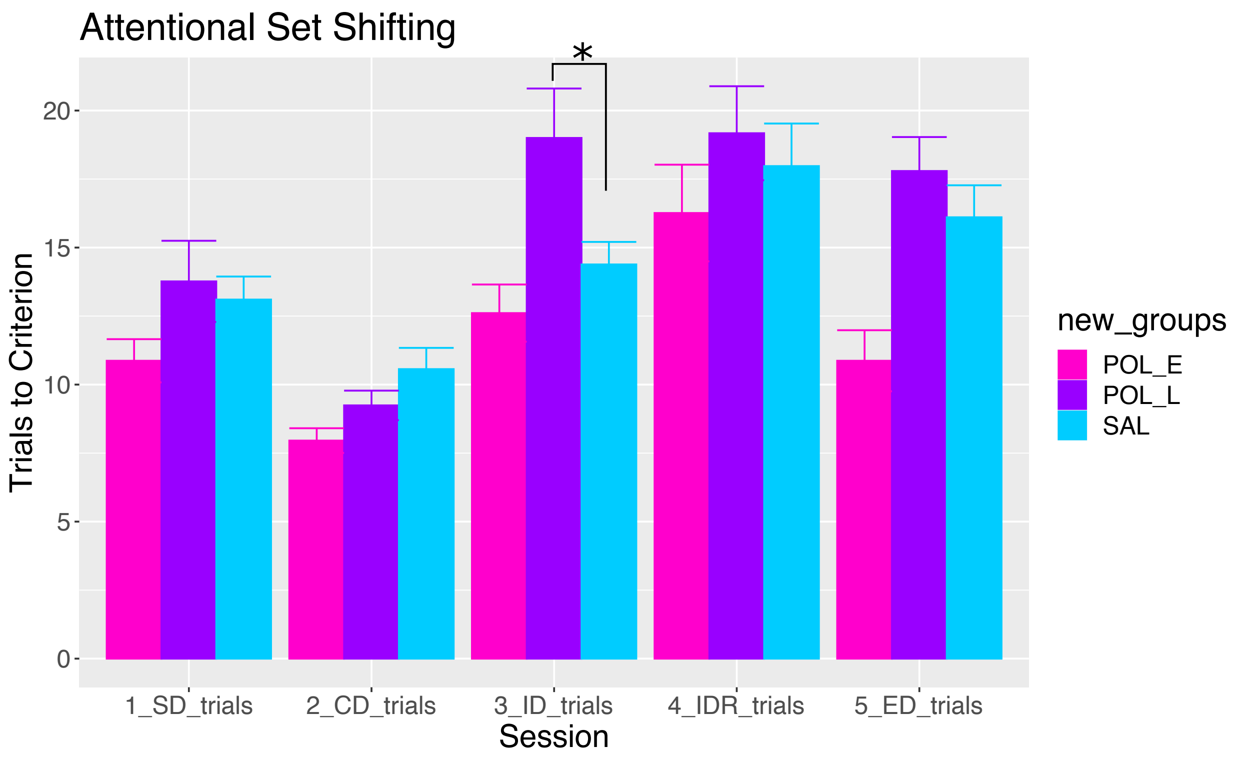


**Supplementary figure 12. Attentional set shifting task results.** Bar graph showing trails to reach learning criterion (6 correct trials in a row) for each portion of the task: **SD=** simple discrimination, **CD=** compound discrimination, **ID=** Intradimensional shift, **IDR=** Intradimensional shift reversal, **ED=** Extra-dimensional shift. Only subtle differences emerged, with POL L requiring more trials in the ID to reach learning criterion (t=2.486, p=0.0129). *p<0.05.

#### 2.7.2. Behavioural sex differences

No sex differences were observed in either the open field task or the marble burying task.

##### 2.7.2.1. Social preference and social novelty

Examination of sex differences in social preference did not result in a sex * group interaction for adolescent mice, however there was a trend level effect for the main effect of POL E treatment (t= -1.893, p=0.0671), wherein POL E offspring, particularly males, had a lower sociability index. In adulthood, we observed a trend level sex by group interaction for POL E offspring (t=-1.829, p=0.0712) wherein POL E males actually had a higher sociability index relative to SAL with no difference for females. The POL E main effect was also significant in this model (t=2.627, p=0.0124). No sex effects were observed in adolescent offspring for social novelty behaviour. The same was true for adult behaviour, however the POL E main effect was significant (t=-2.051p=0.0471).

##### 2.7.2.2. Sensorimotor Gating

We also explored sex differences and observed no singificant sex by group by prepulse level interactions, however this model revealed a number of trend level group by prepulse level interactions for POL E offspring at PP15 (t=-1.830, p=0.0691), and for POL L offspring at PP12 (t=-1.784, p=0.0764). Sex differences were also investigated with increasing prepulse level. There were no sex by group by level interactions in adolescence, however the group by level interaction for POL E (t=-1.755, p=0.0809) and the sex by level interaction (t=-1.957, p=0.0520) both trended towards significance, indicative of the fact that females, and POL E offspring had lower percent PPI. No effects were observed in adulthood.

##### 2.7.2.3. Attentional Set Shifting Task

Exploration of sex differences revealed a number of sex * group * trial interactions as follows. For the CD portion of the task the three way interaction was significant both for POL E (t=2.156, p=0.0311) and POL L (t=2.138, p=0.0325) groups vs. SAL wherein POL E females required more trials than males to reach learning criterion and POL L females required fewer trials than males. Further, the three way interaction was also significant for the POL L group relative to SAL for the IDR (t=2.690, p=0.0071 ) and ED (t=4.140 p=3.47e-05) portions of the task wherein POL L females required more trials to reach learning criterion than males (relative to SAL controls).

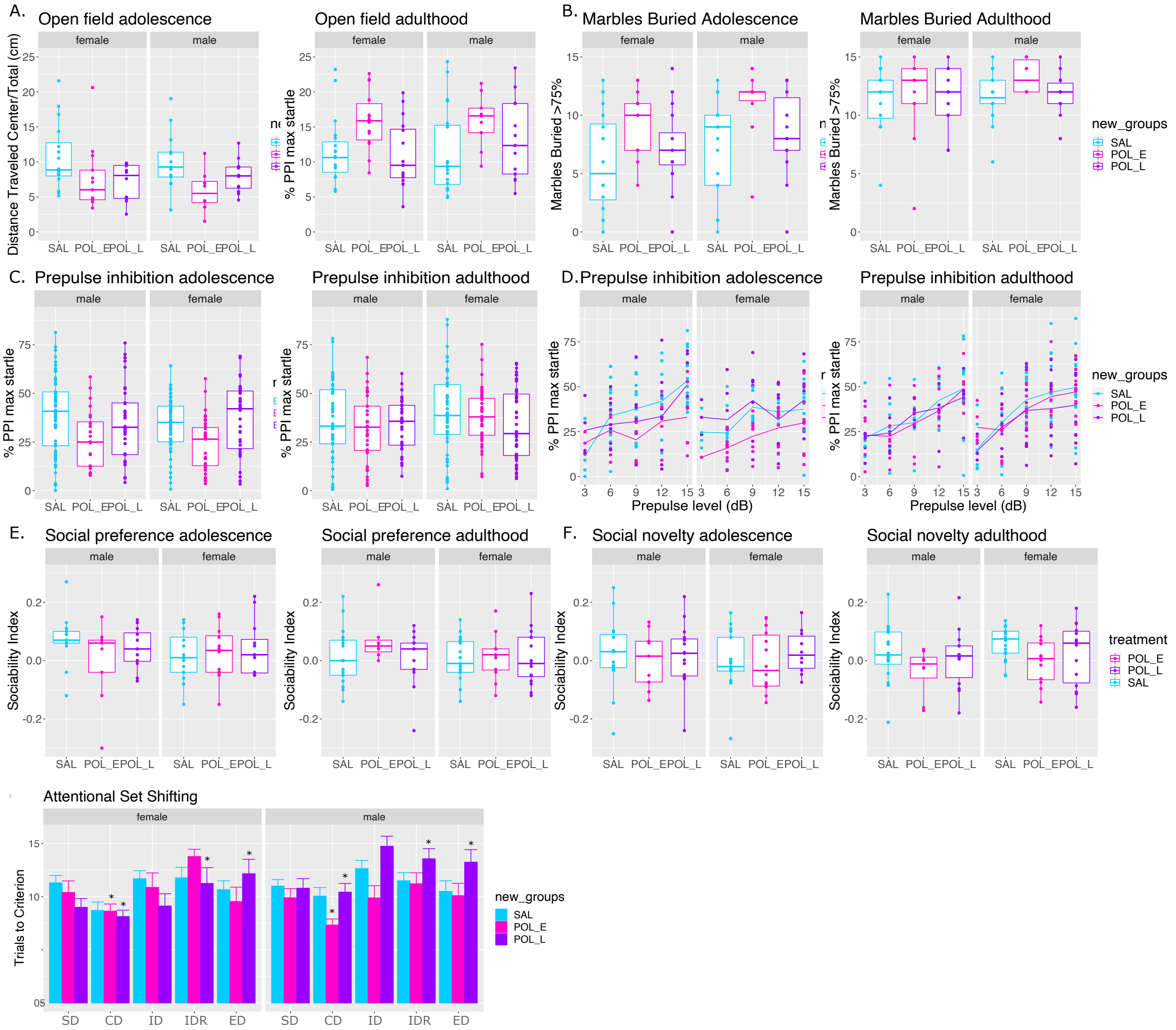


**Supplementary figure 13. Behavioural results split by sex.** Results for adolescent (left) and adult (right) offspring from our three groups: SAL (cyan), POL E (magenta), POL L (purple), with male results on the left, panels, and female results on the right. For all boxplots the midline represents the median of the data, the box represents the first and third quartiles, and the vertical line and dots represent the end range of the data. **(A)** Open field data for distance traveled in the center zone relative to total distance traveled. In adolescence (left), results. No statistically significant differences observed in adulthood (right). (**B**). Significantly more marbles buried by POL E adolescent offspring (left; results and no differences in adulthood (right). (**C**) results. (**D**) results. (**E**) No significant differences in sociability index for the social preference task (i.e. preference for novel mouse over nonsocial object) between groups at either adolescence (left) or adulthood (right). (**F**) No significant differences in sociability index for social novelty (i.e. preference for novel mouse over familiar mouse) between groups. (**F**) ASST results

• $p<0.05$; *$p<0.0045$ (Bonferroni correction threshold); ***$p<0.0001$

### 2.8. Multivariate analysis of brain-behaviour data

The second latent variable (LV2: 19% covariance explained, p=0.002) identified a pattern of brain-behaviour driven by sex and litter size. Being female, and coming from a large litter was associated with smaller volume of regions in blue, such as motor, somatosensory, cingulate, auditory, and visual cortices, nucleus accumbens, amygdala, hippocampus.

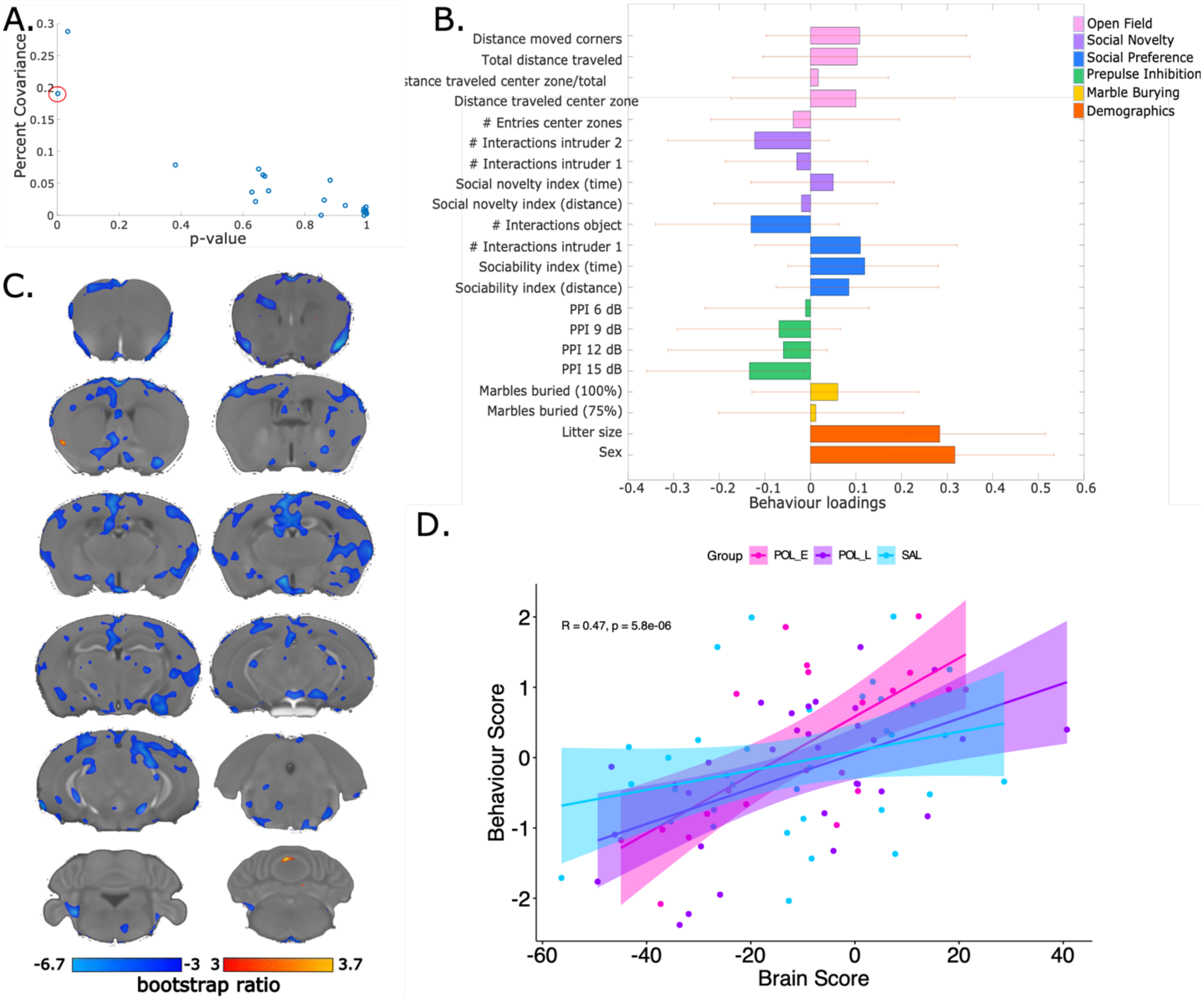


**Supplementary figure 14.** PLS results for LV2. (**A**) Covariance explained (y-axis) and permutation p-values (x-axis) for all 21 LVs in the PLS analysis. LV2 is circled in red (19% covariance explained, p=0.002) (**B**) Behaviour score pattern for each behavioural measure included in the analysis. Singular value decomposition estimates the size of the bars, and confidence intervals are calculated by bootstrapping. (**C**) Brain pattern bootstrap rations overlaid on the population average, with positive bootstrap ratios in orange-yellow, and negative in blue. **(D**) Correlation of individual mouse brain and behaviour score color coded by treatment group.

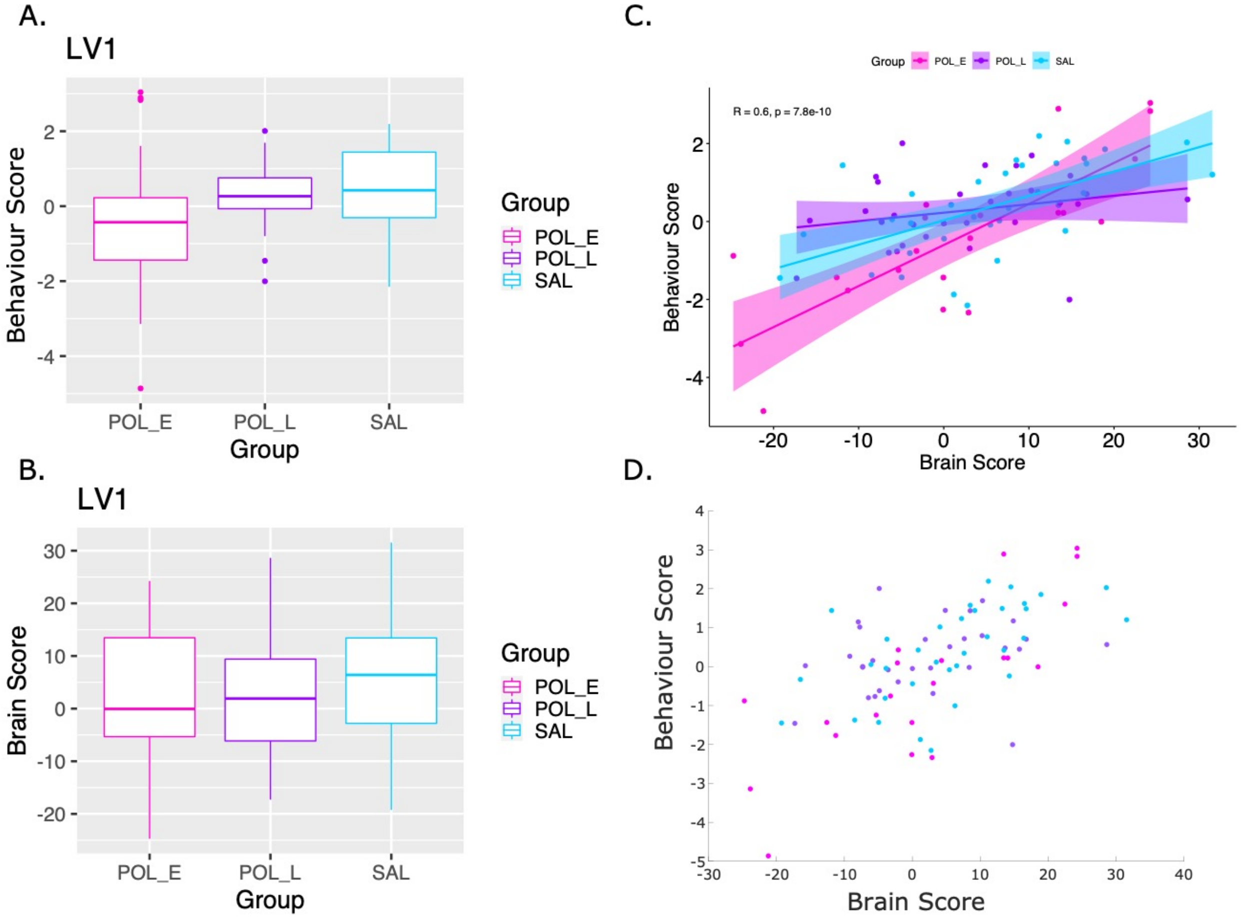


**Supplementary figure 15.** Boxplot for LV1 behaviour score **(A)** and brain score **(B)** for each of the three groups. (**C).** Correlation between brain-behaviour weights in adolescence with a correlation line per group, highlighting that the POL E group has the strongest brain-behaviour correlation for the LV1 pattern. (**D).** Correlation of brain-behaviour without one POL E subject that was an outlier in their behaviour score.

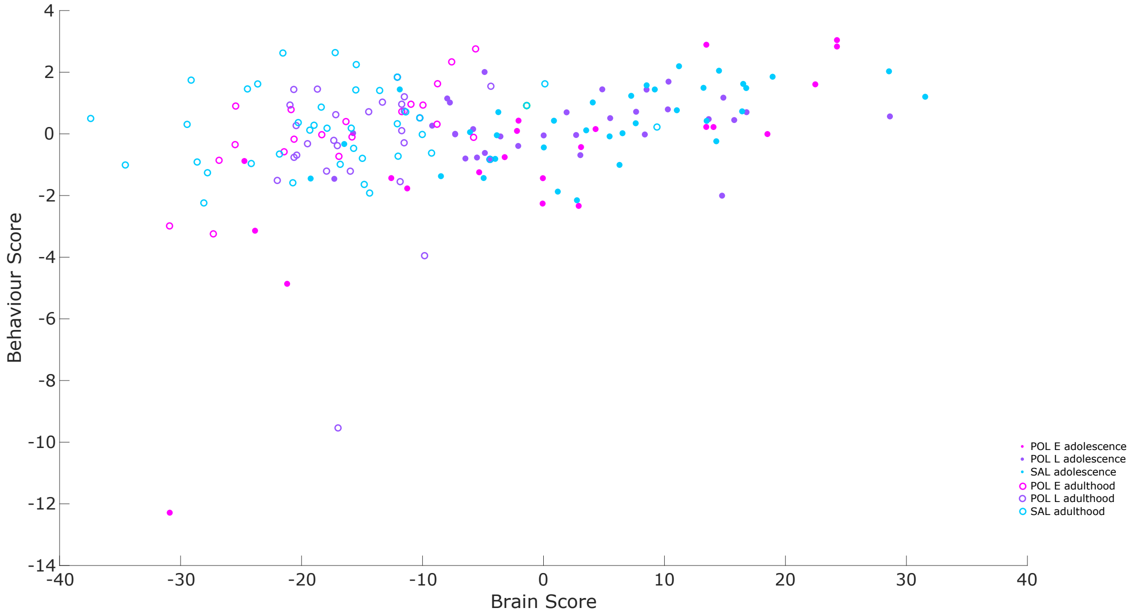


**Supplementary figure 16.** Correlation between the brain-behaviour weights for LV1computed on adolescent data (filled in dots) multiplied by brain and behaviour data for the same tests and same mice in adulthood (open dots). SAL (cyan), POL E (magenta), POL L (purple)

#### 2.9. Transcriptional results

Summary tables for differential gene expression (see appendix):

**Supplementary table 4.** Differential gene expression for POL E vs SAL E across all ROIs (global), ACC, dHIP, and vHIP

**Supplementary table 5.** g:Profiler results for pathway enrichment analysis for POL E vs SAL for all ROIS, ACC, dHIP, and vHIP, for both up- and downregulated DEGs.

**Supplementary table 6**. Gene overlap between published human transcriptional findings for schizophrenia.

##### 2.9.1. Sex differences in differential gene expression

When further breaking down the groups to investigate sex effects, more DEGs were identified in male than female mice in all ROIs. For the dHIP, 199 down- and 381 upregulated genes were observed in males, and 0 in females. For the vHIP 2 down- and 2 upregulated genes were observed in males, whereas 31 down- and 11 upregulated genes were observed in females, and for the ACC 28 down- and 4 upregulated genes were observed in males, and 1 down regulated gene in females.

Upregulated genes for males in the ACC were enriched for non-homologous end joining genes, involved in repairing double-strand breaks in DNA, whereas downregulated genes were enriched for various inflammatory markers including TNF signaling, T cell receptor pathways, IL-17 and IL-1 signaling, as well as apoptosis, and oxidative stress. In the male dHIP, upregulated genes were enriched for synapse structure and function, dendrites, and neurotransmitter release, whereas downregulated genes were enriched for HDMs demethylate histones, and various miRNAs (miR-499-5p, miR-208b-3p, miR-208a-3p). Finally, the male vHIP upregulated genes were enriched for metal sequestration by antimicrobial proteins.

For females, upregulated vHIP genes were enriched for IL-1 signaling, as well as sweat gland function. No other ROIs had significant enchantment results.

**2.9.2. Sex differences in RRHO**

The results were very concordant as expected, we interestly observed the vHIP brain region as carrying most of the differential expression overlap concordance signal between males and females (supplementary figure 17). The strongest overlap for males was for coordinately upregulated genes in the dHIP and vHIP (4878), followed by downregulated genes in these two ROIs (4271). There was also robust overlap for ACC and dHIP upregulated genes (4102), and to a lesser extent for concordantly downregulated genes (2365). Overlap between the ACC and vHIP was also observed with 3252 coordinately upregulated and 3127 downregulated genes. Genes coordinately downregulated in the ACC and dHIP, ACC and vHIP, and dHIP and vHIP were enriched for ribosomal function, translation, and mRNA. Coordinately upregulated genes for all pairs of ROIs were enriched for myelination, oligodendrocyte differentiation, synaptic vesicle regulation, and intracellular transport.

For females, the strongest overlap was observed for coordinately downregulated genes in the ACC and vHIP (4116), followed by upregulated genes in the ACC and dHIP (3800), upregulated genes in the ACC and vHIP (3615), and the dHIP and vHIP (3186). Surprisingly there was very little overlap in concordantly downregulated genes in the ACC and dHIP (830) and dHIP and vHIP (247). Pathway analysis revealed concordantly upregulated genes in all pairs of ROIs were enriched for mitochondrial function, translation, and ribosomal function, in addition to synaptic function for the dHIP and vHIP upregulated genes. Concordantly downregulated genes were less homogeneous; in the ACC and dHIP overlap, enrichment was observed for protein-protein interactions at synapse; for the ACC and vHIP, enrichment was observed for cilium assembly and organization, extracellular structural matrix, microglia pathogen phagocytosis pathway; finally, for the dHIP and vHIP, enrichment was observed for serotonin and anxiety related events, complement and coagulation cascades, IL-17 signaling.

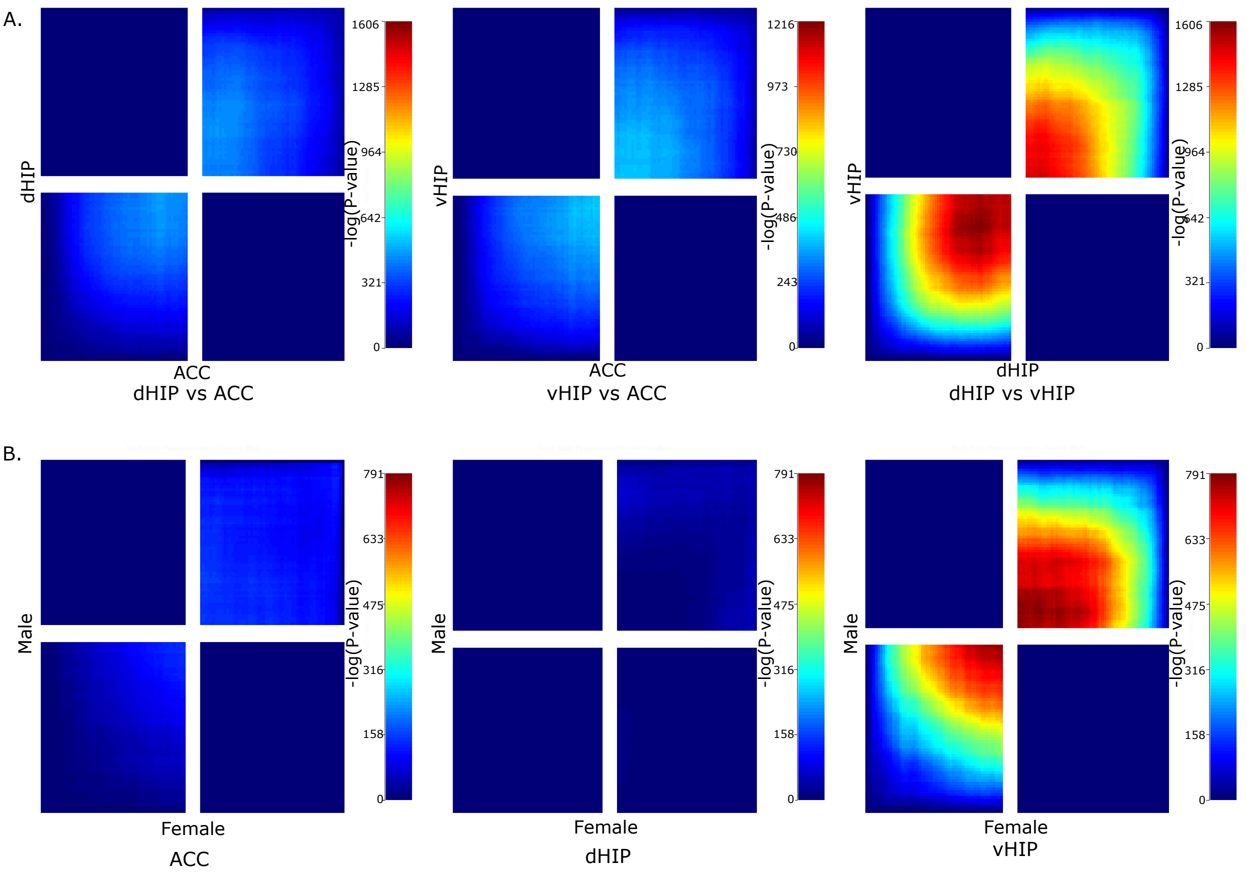


**Supplementary figure 17. A**. RRHO heatmaps for the dHIP vs ACC (left), vHIP vs ACC (middle) and vHIP vs dHIP (right). For heat maps, the top left quadrant indicates the overlap in genes up-regulated in the first region and downregulated in the second region. The top right quadrant indicates overlap in genes downregulated in both regions. The bottom left quadrant indicates overlap in genes upregulated in both regions. The bottom right quadrant indicates overlap in genes downregulated in the first region and upregulated in the second. **B**. RRHOs heatmaps comparing up- and downregulated genes in males vs. females for each ROI, with ACC (left), dHIP (middle), vHIP (right).

Friedel, Miriam, Matthijs C. van Eede, Jon Pipitone, M. Mallar Chakravarty, and Jason P. Lerch. 2014. “Pydpiper: A Flexible Toolkit for Constructing Novel Registration Pipelines.” *Frontiers in Neuroinformatics* 8 (July): 67.

Fuentes, Montserrat, Peter Guttorp, and Paul Sampson. 2006. “Using Transforms to Analyze Space-Time Processes.” *C&H/CRC Monographs on Statistics & Applied Probability*. https://doi.org/10.1201/9781420011050.ch3.

Kovačević, N., J. T. Henderson, E. Chan, N. Lifshitz, J. Bishop, A. C. Evans, R. M. Henkelman, and X. J. Chen. 2004. “A Three-Dimensional MRI Atlas of the Mouse Brain with Estimates of the Average and Variability.” *Cerebral Cortex* 15 (5): 639–45.

Lerch, Jason P., Jeffrey B. Carroll, Shoshana Spring, Lisa N. Bertram, Claudia Schwab, Michael R. Hayden, and R. Mark Henkelman. 2008. “Automated Deformation Analysis in the YAC128 Huntington Disease Mouse Model.” *NeuroImage* 39 (1): 32–39.

Meyer, Urs, Myriel Nyffeler, Andrea Engler, Adrian Urwyler, Manfred Schedlowski, Irene Knuesel, Benjamin K. Yee, and Joram Feldon. 2006. “The Time of Prenatal Immune Challenge

Determines the Specificity of Inflammation-Mediated Brain and Behavioral Pathology." *The Journal of Neuroscience: The Official Journal of the Society for Neuroscience* 26 (18): 4752–62.
Patel, Raihaan, Christopher J. Steele, Anthony G. X. Chen, Sejal Patel, Gabriel A. Devenyi, Jürgen Germann, Christine L. Tardif, and M. Mallar Chakravarty. 2020. "Investigating Microstructural Variation in the Human Hippocampus Using Non-Negative Matrix Factorization." *NeuroImage* 207 (February): 116348.
Ritchie, Matthew E., Belinda Phipson, Di Wu, Yifang Hu, Charity W. Law, Wei Shi, and Gordon K. Smyth. 2015. "Limma Powers Differential Expression Analyses for RNA-Sequencing and Microarray Studies." *Nucleic Acids Research* 43 (7): e47.
Robinson, Mark D., and Alicia Oshlack. 2010. "A Scaling Normalization Method for Differential Expression Analysis of RNA-Seq Data." *Genome Biology* 11 (3): R25.
Semple, Bridgette D., Klas Blomgren, Kayleen Gimlin, Donna M. Ferriero, and Linda J. Noble-Haeusslein. 2013. "Brain Development in Rodents and Humans: Identifying Benchmarks of Maturation and Vulnerability to Injury across Species." *Progress in Neurobiology*. https://doi.org/10.1016/j.pneurobio.2013.04.001.
Tustison, Nicholas J., Brian B. Avants, Philip A. Cook, and James C. Gee. 2010. "N4ITK: Improved N3 Bias Correction with Robust B-Spline Approximation." *2010 IEEE International Symposium on Biomedical Imaging: From Nano to Macro*. https://doi.org/10.1109/isbi.2010.5490078.
Zeighami, Yashar, Seyed-Mohammad Fereshtehnejad, Mahsa Dadar, D. Louis Collins, Ronald B. Postuma, Bratislav Mišić, and Alain Dagher. 2019. "A Clinical-Anatomical Signature of Parkinson's Disease Identified with Partial Least Squares and Magnetic Resonance Imaging." *NeuroImage* 190 (April): 69–78.
